# Directed Chemical Evolution via Navigating Molecular Encoding Space

**DOI:** 10.1101/2025.03.18.643899

**Authors:** Lin Wang, Yifan Wu, Hao Luo, Minglong Liang, Yihang Zhou, Cheng Chen, Chris Liu, Jun Zhang, Yang Zhang

## Abstract

Deep-learning techniques have significantly advanced small-molecule drug discovery. However, a critical gap remains between representation learning and small molecule generations, limiting their effectiveness in developing new drugs. We introduce Ouroboros, a unified framework that integrates molecular representation learning with generative modeling, enabling efficient chemical space exploration using pre-trained molecular encodings. By reframing molecular generation as a process of encoding space compression and decompression, Ouroboros resolves the challenges associated with iterative molecular optimization and facilitates directed chemical evolution within the encoding space. Comprehensive experimental tests demonstrate that Ouroboros significantly outperforms conventional approaches across multiple drug discovery tasks, including ligand-based virtual screening, chemical property prediction, and multi-target inhibitor design and optimization. Unlike task-specific models in traditional approaches, Ouroboros leverages a unified framework to achieve superior performance across diverse applications. Ouroboros offers a novel and highly scalable protocol for rapid chemical space exploration, fostering a potential paradigm shift in AI-driven drug discovery.

## Introduction

Artificial neural networks have shown remarkable ability to autonomously identify underlying data structures through their advanced representation learning capabilities^1,2^. Many techniques originally developed for natural language processing and image generation^3–5^ have been successfully applied to drug discovery. For instance, considerable progress has been recently achieved in leveraging deep neural networks to learn interpretable molecular representations and encodings^6^, which can be of critical help in molecular property prediction and ligand-based virtual screening^7–11^.

Most recently, growing interests have been shown in generative AI for drug design. This approach often builds upon Quantitative Structure-Activity Relationship (QSAR), a technique originally developed 60 years ago^12^ and now enhanced by molecular representation learning models to evaluate molecules produced by molecular generators^13^. While it helps facilitate iterative molecular generation, it limits models’ capacity to perceive optimization directions. To partly address this, researchers have explored integrating drug property prediction and binding pocket condition into the training process of generative models^14–18^. However, this integration has posed considerable challenges in model extrapolation and reusability, mainly due to the inherent disconnections between molecular representation learning and molecular generation, a critical yet often overlooked gap that impedes the direct application of insights gained from representation learning into generative models.

Another emerging approach to molecular generation involves searching for neighbors of known drug molecules within the chemical space^19^. The effectiveness of the approach is highly correlated with the relationship between chemical space and encoding space learned by representation models. Although evidence suggests that chemical structures and neural network encodings may share common grounds, e.g., similar chemical structures often exhibit analogous biological activities, just as similar encodings yield comparable property outputs, a robust model for quantitatively mapping encoding to chemical space remains absent, hindering the effective application of the neighbor-search-based generative models in drug discovery.

To address these challenges, we introduce Ouroboros (as shown in **Figure 1a**), a new protocol to bridge the gap between representation learning and generative AI models and facilitate chemical space navigation for drug discovery. For doing this, Ouroboros first employs representation learning to encode molecular graphs into 1D vectors. These encodings are then independently decoded to predict both molecular properties and molecular structures, a process also known as chemical space decompression. One aim of Ouroboros is to transform the focus from traditional virtual screening^9^, which often relies on property predictors and encoding similarities for screening compound library, to directed chemical evolution within the encoding space to improve the effectiveness (**Figure S1**). Additionally, leveraging the encoding space for molecular structure decompression enables generative AI to integrate chemical knowledge from representation learning for guiding the directed chemical evolution of molecules. To examine the practical feasibility, we present a multi-level case study of the Ouroboros protocol, involving an independent representation learning for molecular encoder (**Figure 1b**), a molecular structure decoder to reconstruct the molecular encoding to molecular SMILES (**Figure 1c**), and a molecular property decoder to navigate the directed chemical evolution (**Figure 1d**). Furthermore, we explore three distinct strategies for chemical evolution, offering insights into the practical applications of Ouroboros in drug discovery.

**Figure 1.**
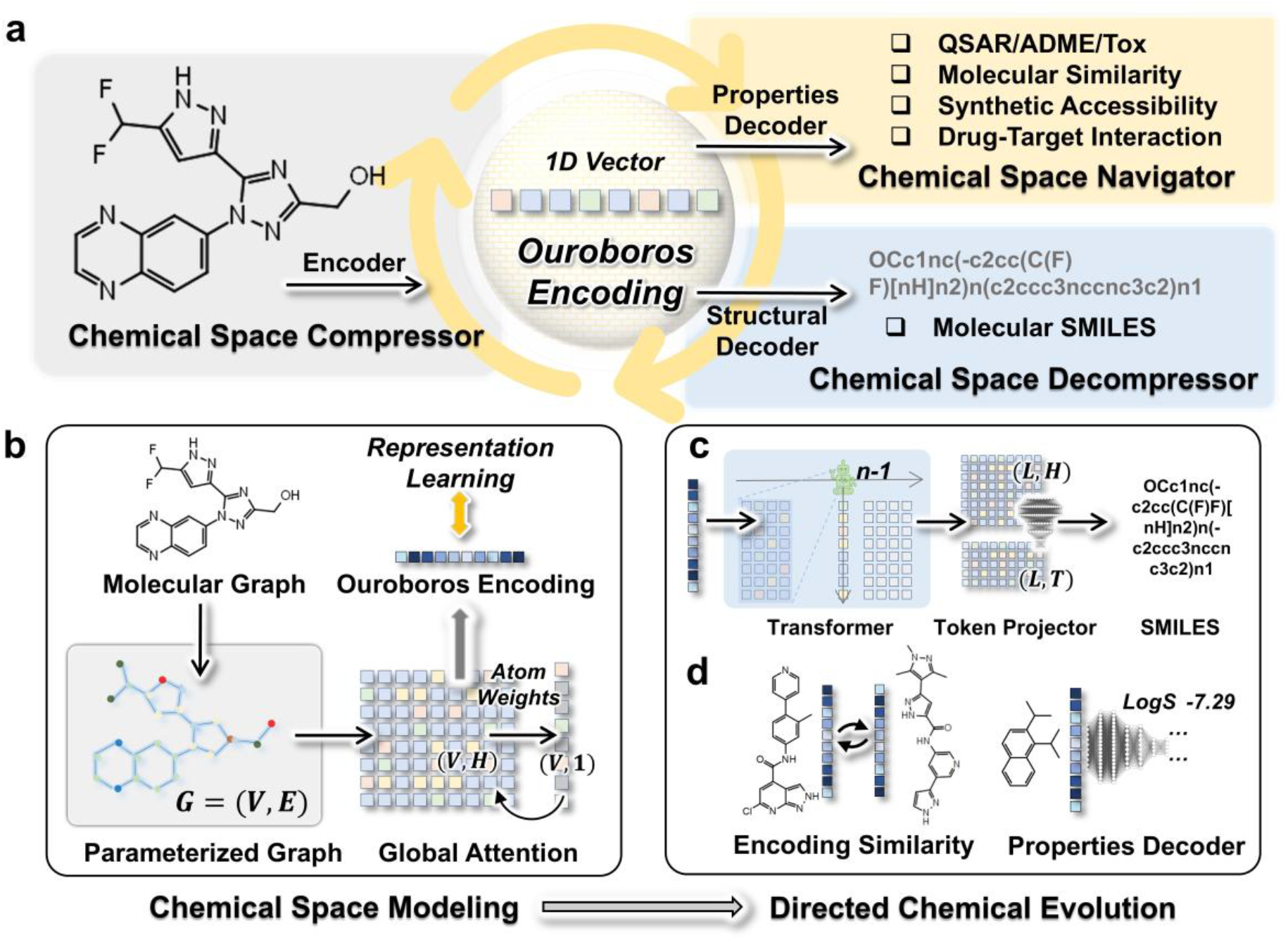
The Ouroboros protocol for chemical space navigation. **(a)** Three independent modules of Ouroboros. The chemical space compressor converts molecular graphs into 1D vectors, typically implemented by a graph neural network (GNN), while the property decoder projects the 1D vectors onto molecular properties through a multilayer perceptron (MLP). Finally, the chemical space decompressor transforms these 1D vectors into SMILES representations, utilizing a text generation model by Transformer decoder. **(b)** The molecular encoder for chemical space modeling, where the molecular graph is encoded by a global-attention based GNN module and represented as a 1D encoded vector, with ‘E’ referring to bonds, ‘V’ to atoms, and “H” to hidden size. **(c)** The molecular decoder for decompressing the molecular encoding to molecular structure, with ‘L’ refer to sequence length. **(d)** The two approachs for chemical space navigations.

## Results

### Compressing the chemical space through similarity learning

Representation learning creates a structured encoding space to interpret chemical molecular structures, with the assumption that molecules with similar chemical structures or pharmacological properties are positioned in proximity within encoding space. In a previous study, we demonstrated that incorporating Conformational Space Similarity (CSS) yields a generic molecular representation for ligand-based drug discovery^9^. To improve the performance of molecular representation learning, here we introduce two kinds of inter-molecular similarities, including an extended CSS with enhanced space searching and molecular fingerprint similarity (MFS) (see **Methods**). Furthermore, we propose a new similarity learning strategy that projects molecular encodings into multiple molecular similarities, aiming to enhance the capacity of the molecular encoder for generic and informative molecular encoding.

The pre-training architecture of representation learning in Ouroboros is outlined in **Figure S2**. The molecular encoder, which transforms molecular graphs into 1D encoding vectors, is pre-trained on a similarity matrix constructed from pairwise similarities between query and reference molecules. The 1D representations are then projected by two external projection heads into MFS and CSS respectively with the training process guided by the Mean Squared Error (MSE) loss. As shown in **Figure S3**, the Spearman correlation on the validation dataset reached convergence after about 20,000 pre-training steps and showed a similar performance on the test set (**Table S1**).

**Figure 1b** illustrates how the molecular encoding is extracted from molecule graphs, where atoms and chemical bonds in the molecular graphs are represented through a GNN for message passing. The resulting parameterized graph is then processed through a global attention pooling module^20^ to construct the molecular encoding. The pre-trained encoder captures the varying contributions of different atoms to the overall molecular encoding. For atoms of the same element type but with distinct pharmacophore features, the Ouroboros encoder assigns differentiated weights. For example, a nitrogen atom with positive charge is weighted differently from those in amide groups (**Figure S4**). Moreover, the model demonstrates heightened attention to bulky hydrophobic groups when they appear, emphasizing their importance in pharmacophore and conformational flexibility.

### Benchmarking the Ouroboros encoder for virtual screening and property prediction

Directed chemical migration within the Ouroboros encoding space requires diverse property decoders to guide the optimization process, such as molecular similarity and molecular property predictors (**Figure S1**). Consequently, the generality and versatility of molecular encodings play a critical role in determining the capability of Ouroboros in drug discovery. Ideally, these encodings should be trained on a limited subset of molecular structures while maintaining the ability to generalize across large-scale benchmark datasets, such as 1,463,336 molecules in DUD-E^21^ and 2,808,885 in LIT-PCBA^22^. Additionally, they should exhibit superior performance across various molecular property modeling tasks to ensure broad applicability and robustness.

#### Ligand-based virtual screening

To examine the potential of Ouroboros encoder for virtual screening, we estimate the chemical similarity of two molecules based on the cosine or Pearson similarity between 1D encoding vectors. **Figure 2a** compares it with 6 baseline methods using an early enrichment metric called Boltzmann-enhanced discrimination of receiver operating characteristic (BEDROC)^23^ across two virtual screening benchmarks (DUD-E^21^ and LIT-PCBA^22^). The results show that despite the relatively small training dataset, Ouroboros encoding achieves the highest enrichment scores in both benchmarks. Furthermore, as shown in **Table S2**, Ouroboros outperformed the baseline methods on most of other metrics, including AUPRC, AUROC, enrichment factor (EF), and log AUC. These results suggest that the Ouroboros encodings are more effective on clustering molecules with similar pharmacological features in proximity within the encoding space compared to the baseline methods, supporting the use of Ouroboros encoding similarity in directed chemical migration.

**Figure 2.**
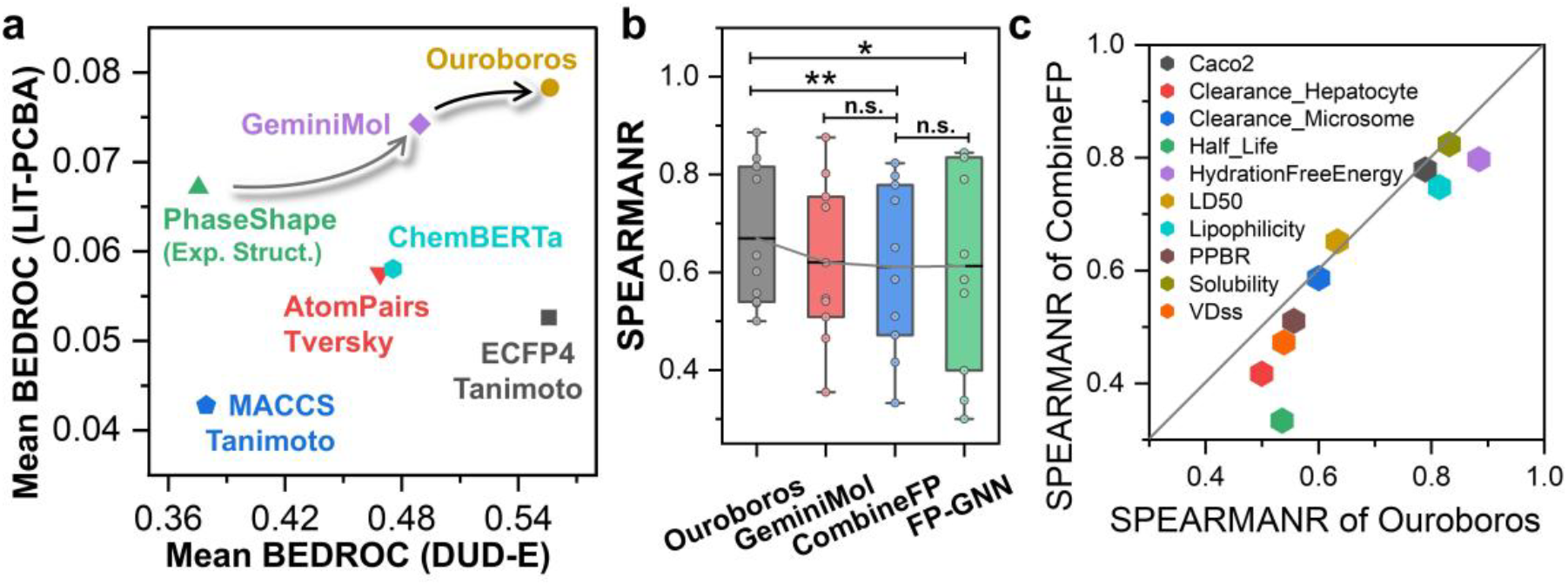
Assessing the quality of chemical space modeling in Ouroboros. **(a)** Similarity-based virtual screening performance evaluated by the mean BEDROC on the 102 targets from DUD-E versus that on the 15 targets from LIT-PCBA by different methods, including Ouroboros, GeminoMol, PhaseShape, ChemBERTa (the best performend version of ChemMLM), AtomPairs, ECFP4, and MACCS. **(b)** Spearman correlation coefficients achieved by three molecular encoders on 10 different molecular property regression tasks. For GeminiMol and Ouroboros, the decoder is constructed using a multilayer perceptron. CombineFP combines four molecular fingerprints together, including AtomPairs, TopologicalTorsion, ECFP4, and FCFP6, which uses AutoGluon to construct property predictors. ‘^**^’ refers to p-value=0.0079, ‘^*^’ to p-value=0.0540, and ‘n.s’ to not significant (p-value>0.3). **(c)** Spearman correlation coefficients by Ouroboros on 10 different molecular property regression tasks (colored differently) versus that by CombineFP.

#### Chemical property prediction

The molecular property predictor in Ouroboros is implemented by mapping molecular encodings to properties via deep neural networks. **Figure 2b** presents the average Spearman correlation coefficient (SPEARMANR) of the Ouroboros predictor on 10 different molecular property datasets, which is significantly higher than that of AI-based GeminiMol and fingerprint-based CombineFP^9^. It also outperformed FP-GNN^24^, which was specifically designed for molecular property modeling by integrating multiple fingerprints with GNNs. If we check the individual properties, Ouroboros achieved the highest SPEARMANR in six out of 10 property tasks among all four predictors (**Table S3**). As an illustration, **Figure 2c** shows a head-to-head comparison of Ouroboros with CombineFP, where Ouroboros not only excels in relatively simple tasks, such as lipophilicity and hydration free energy prediction, but also achieves substantial improvements in more challenging tasks like steady-state volume of distribution and clearance rate. These results underscore the strong generalizability of Ouroboros encodings, which are crucial for training effective property predictors across a diverse range of downstream molecular property tasks.

### Multi-target cancer inhibitor discovery using Ouroboros molecular encoder

#### Strategy for multi-target cancer inhibitor identification

Similarity-based virtual screening is a widely used approach in ligand-based drug discovery, aiming to identify novel chemical scaffolds by retrieving new molecules that resemble known active compounds. Given the outstanding performance of Ouroboros in the DUD-E and LIT-PCBA benchmarks, we apply the encoder to a challenging multi-target drug discovery task to evaluate its practical utility. This task focuses on identifying multi-target inhibitors for five common cancer driver gene mutations, targeting synthetic lethality (for tumor suppressor genes) or promoting cell proliferation (for oncogenes).

A total of 10 distinct drug targets and 119 reference compounds are included in the experiment. Following the strategy outlined in **Figure 3a**, we enrich potential active molecules through similarity screening, followed by molecular docking to select candidate compounds for experimental assay. We anticipated that Ouroboros would identify structurally novel inhibitors from the Enamine REAL diversity set^25^, capable of simultaneously targeting at least two of these drug targets.

**Figure 3.**
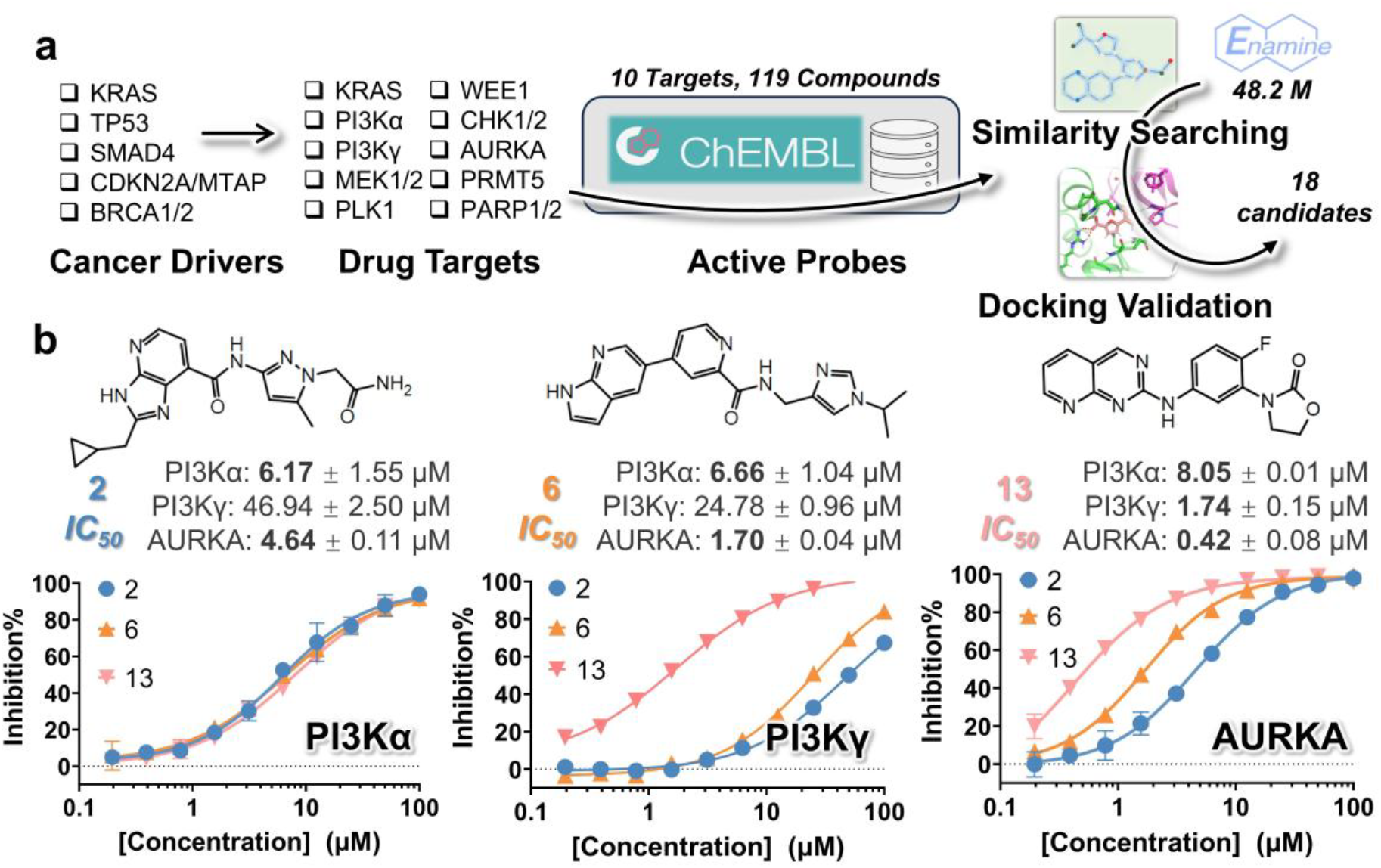
The dicovery of novel multi-target inhibtors using Ouroboros. **(a)** A pipeline for screening potential multi-target cancer driver inhibitors. **(b)** *IC50* and inhibition curves of three hit compounds (#2, #6 and #13) for three targets, including PI3Kα, PI3Kγ and AURKA.

#### Experimental validation

A total of 18 molecules are successfully synthesized for enzyme activity assay, of which 7 compounds (38%) are active, and 3 compounds (16%) exhibited multi-target inhibitory behavior as expected (**Table S4**). **Figure 3b** summarizes the results of the 3 identified hit molecules, each achieving a maximum similarity of over 0.5 with the active reference molecules of three targets of interest, including PI3Kα, the phosphatidylinositol 4,5-bisphosphate 3-kinase catalytic subunit alpha isoform; PI3Kγ, the phosphatidylinositol 4,5-bisphosphate 3-kinase catalytic subunit gamma isoform; and AURKA, the aurora kinase A, which demonstrates appreciable inhibitory activity in enzymatic assays.

Notably, compound #13 exhibited superior activity across all three targets, with an *IC50* in the nanomolar range for AURKA, highlighting its potential as a promising lead compound. However, the testing results of all 18 candidate compounds (**Table S4**) indicated that none showed an ability to inhibit more than four targets simultaneously. This limitation may be attributed to the relatively low throughput of the experiment and the use of only a 48.2M subset of the REAL library^25^ during the screening. Additionally, the findings confirmed that compounds #2, #6, and #13 are not pan-kinase inhibitors, which selectively inhibit PI3K and AURKA. Overall, we identified three multi-target hit molecules from a 48.2M compound dataset by Ouroboros combined with molecular docking. These results show the application potential of Ouroboros encoding in similarity screening and further corroborate the ability of Ouroboros molecular encoding to generalize to ultra-large molecular structure datasets. This provides credibility to preform directed chemical evolution in this molecular encoding space.

### Decompressing chemical structures from molecular encoding space

Chemical language encoders and decoders are typically trained jointly to ensure the accurate reconstruction of molecular encodings from hidden layers. However, this coupling presents challenges for applying molecular representation learning in generative AI. One key issue is that the information embedded within the intermediate hidden layers of encoder-decoder models often fails to generalize effectively across diverse molecular properties. Furthermore, as shown in **Table S2**, chemical language models such as ChemBERTa^26,27^, do not demonstrate a substantial advantage over traditional molecular fingerprint-based methods in identifying structurally similar molecules with comparable biological activity. These findings indicate that molecular encodings derived from conventional chemical language pre-training strategies may be insufficiently optimized for capturing the chemical space relevant to drug discovery applications.

#### More than 80% structures are rapidly recovered in validation

In Ouroboros, we developed a new molecular structure decoder that reconstructs the original SMILES representations of molecules from their encoded 1D vectors using a Transformer-based architecture (**Figure S5**). As shown in **Figure S6**, the model converges rapidly within a single training epoch. To assess its performance, **Figure S7** presents the AtomPairs^28^ MFS between the actual molecular structures and those reconstructed by the Ouroboros decoder in the validation set, where the mean similarity curve quickly stabilizes at an 80% threshold as batch numbers increase, demonstrating the efficiency and robustness of the Ouroboros decoder in molecular structure recovery from 1D encodings.

#### Similarity declines more rapidly than validity upon perturbations

To further assess the Ouroboros decoder’s ability to explore latent molecular structures within the encoding space, we evaluate whether the model can generate valid and diverse molecular structures by introducing stochasticity into the decoding process. As shown in **Figure S8**, we employed two approaches for randomized decoding: (1) perturbing molecular encoding vectors by adding noise based on the encoding space distribution and (2) introducing randomness into the token sampling process using the Gumbel-max technique^29^. As expected, the similarity between the original and decoded molecules decreases with increasing temperature and noise levels. However, the similarity curve declines more rapidly than the validity curve (which measures the proportion of decoded molecules with chemically reasonable valence bonds), indicating that Ouroboros effectively generates novel and diverse molecular structures while maintaining structural validity. These findings highlight again the effectiveness of the proposed molecular decoder framework in accurately reconstructing molecular structures from their 1D encodings.

### Directed chemical evolution in encoding space for novel drug generation

One primary objective of Ouroboros is to generate novel and chemically diverse molecular structures beyond the conventional scope of medicinal chemistry. To accomplish this, Ouroboros employs a directed chemical evolution approach, initiating the process with either Gaussian noise or a predefined molecule. The optimization is guided by a loss function constructed from molecular properties, refining the encoding vectors accordingly. By iteratively decoding molecules throughout the optimization, a progressive trajectory of molecular structure evolution emerges, facilitating the discovery of diverse and innovative scaffolds. The resulting molecules can then be selected for further experimental validation, guided by molecular docking, retrosynthetic analysis, and expert input from medicinal chemists (**Figure S9**).

#### Chemical structure propagation

To evaluate the effectiveness of directed chemical evolution, we first examine whether stochastic propagation can progressively modify the structure of the starting molecule and generate novel chemical scaffolds. Using aspirin as an illustrative example, **Figure 4a** demonstrates that the molecular structures generated through this process gradually diverge from the initial molecule while maintaining an observable similarity. Meanwhile, their encoding similarity decreases as propagation progresses. This finding suggests that exploring the neighboring regions of Ouroboros in the encoding space can uncover structural analogs of the starting molecule, supporting the use of Ouroboros for molecular generation.

**Figure 4.**
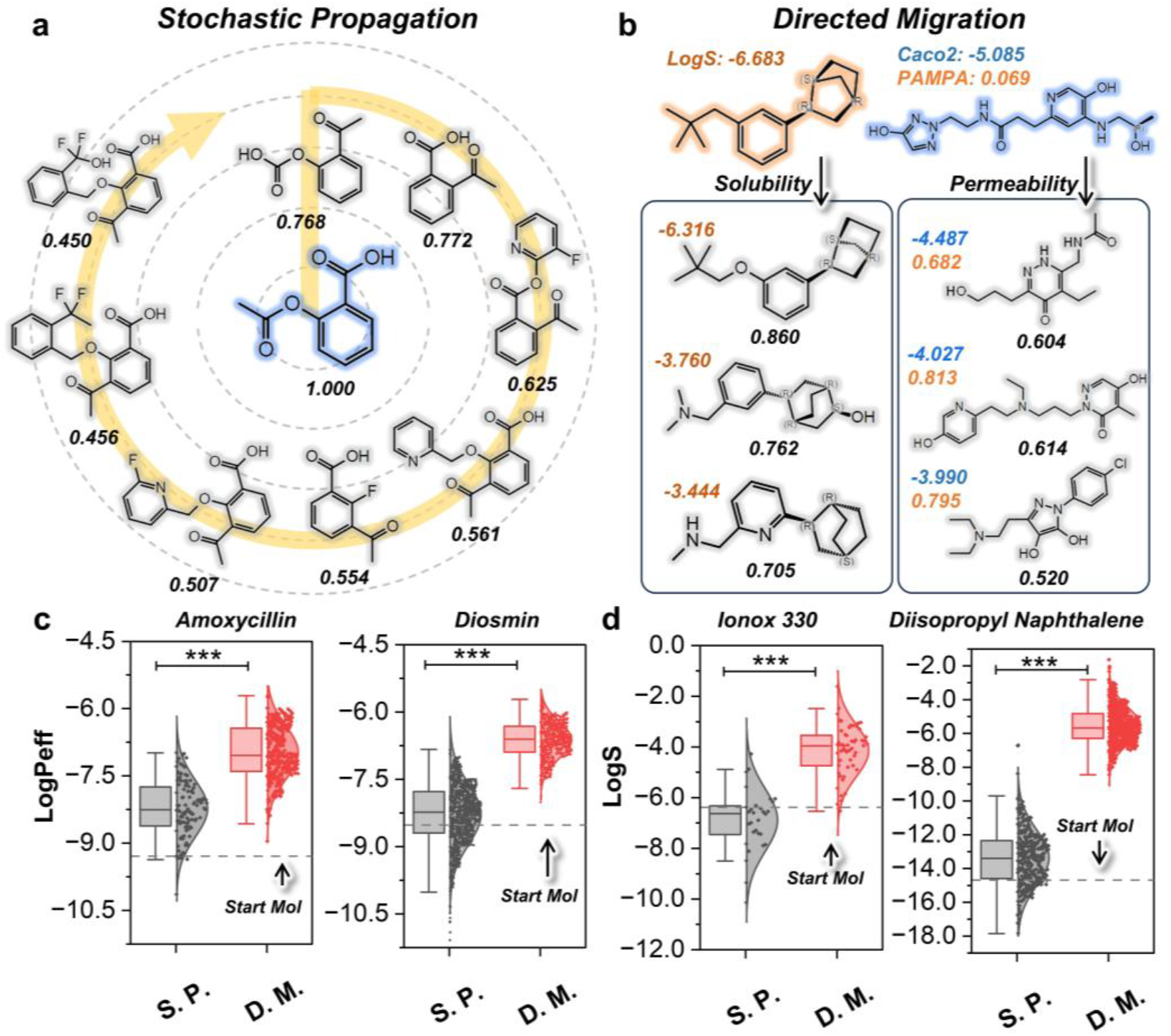
Exploring chemical space with Ouroboros. **(a)** Stochastic propagation from aspirin. Encoding similarities with start molecule are marked at the bottom of molecules. **(b)** Directed migration for optimizing solubility and membrane permeability. The property labels are predictions from the property predictor trained during the benchmark test. The encoding similarity values are displayed beneath each molecule. Solubility is represented by LogS (shown in brown, positioned to the left of each molecule), while membrane permeability is indicated by LogPeff (measured in 10^−6^ cm/s). Parallel artificial membrane permeability values are highlighted in orange, and Caco-2 cell permeability values are shown in blue. PAMPA (Parallel Artificial Membrane Permeability Assay) is also included as a reference. **(c, d)** The comparison of stochastic propagation (S.P.) and directed migration (D.M.) in multi-objective molecular properties optimization for membrane permeability (**c**) and solubility (**d**). The dashed line represents the starting molecule. All the points in the box plots represent new molecules with encoding similarity greater than 0.6 generated during the propagation process and migration pathway. “^***^” refers to p-value < 0.0001.

#### New drug design through directed migration

The opposite of stochastic propagation is directed migration, a strategy that employs a loss function to guide the directed optimization of molecular encoding, thus permitting the directed optimization of chemical structures. As a demo, we examine directed molecular optimization guided by two specific properties: water solubility and membrane permeability. As shown in **Figure 4b**, starting from a hydrophobic molecule, Ouroboros successfully generates molecules with improved solubility while preserving overall structural similarity. Similarly, beginning with a highly flexible and strongly hydrophilic (led to poorly permeability) molecule, Ouroboros produced structures with reduced flexibility and carrying positive charges, which exhibited enhanced predicted membrane permeability. This result demonstrates that Ouroboros encoder is capable of selectively optimizing specific properties of the molecules during the migrations.

To further evaluate the model’s generalizability, **Figures 4c** and **4d** present solubility and membrane permeability optimization results on a broader scale, featuring four representative organic molecules: two with low membrane permeability (Amoxicillin, an antibiotic, and Diosmin, a natural product) and two hydrophobic molecules with poor water solubility (initial structures shown in **Figure S10**). The loss function was designed to simultaneously enhance solubility, membrane permeability, and lipophilicity, while maintaining structural similarity to the original molecules (as described in Methods). Despite applying the same loss function to both categories of molecules with suboptimal properties, Ouroboros’ directed migration successfully generated multiple molecules with notable improvements in membrane permeability or solubility, while preserving an encoding similarity above 0.6. These findings underscore the versatility and effectiveness of Ouroboros’ directed migration approach in molecular property optimization.

#### Directed migration enhances docking scores of chemical compounds

In **Figure S11**, Ouroboros is utilized to map a migration pathway between the encoding vectors of two inhibitors with distinct scaffolds ([1,3,5]-triazine derivative and 5-heterocycle pyrazolo pyridine), both targeting 3’,5’-cyclic-AMP phosphodiesterase 4B (PDE4B), a crucial regulator of various physiological processes^30^. Docking the molecular decoys generated along this pathway into the PDE4B binding pocket revealed that the intermediates retained similar binding poses to the two reference molecules. Notably, three intermediate molecules along the migration trajectory exhibited superior docking scores compared to both reference compounds, highlighting Ouroboros’ potential in scaffold hopping and structural optimization.

### Dual-target drug optimization through decoding-based chemical fusion

The data in **Figure 3b** indicate that AURKA and PI3Kγ are kinase targets amendable for simultaneous inhibition. This finding motivated us to explore whether Ouroboros could integrate pharmacophore features from two sets of reference molecules to generate novel dual-target inhibitors. To investigate this, we conducted similarity screening on the Enamine REAL diversity set^25^ (the training dataset for the molecular decoder) to identify potential dual-target inhibitors for AURKA and PI3Kγ. In parallel, the same reference compounds were input into Ouroboros for chemical fusion, enabling a direct comparison between the candidate compounds generated by both approaches (**Figure 5a**).

**Figure 5.**
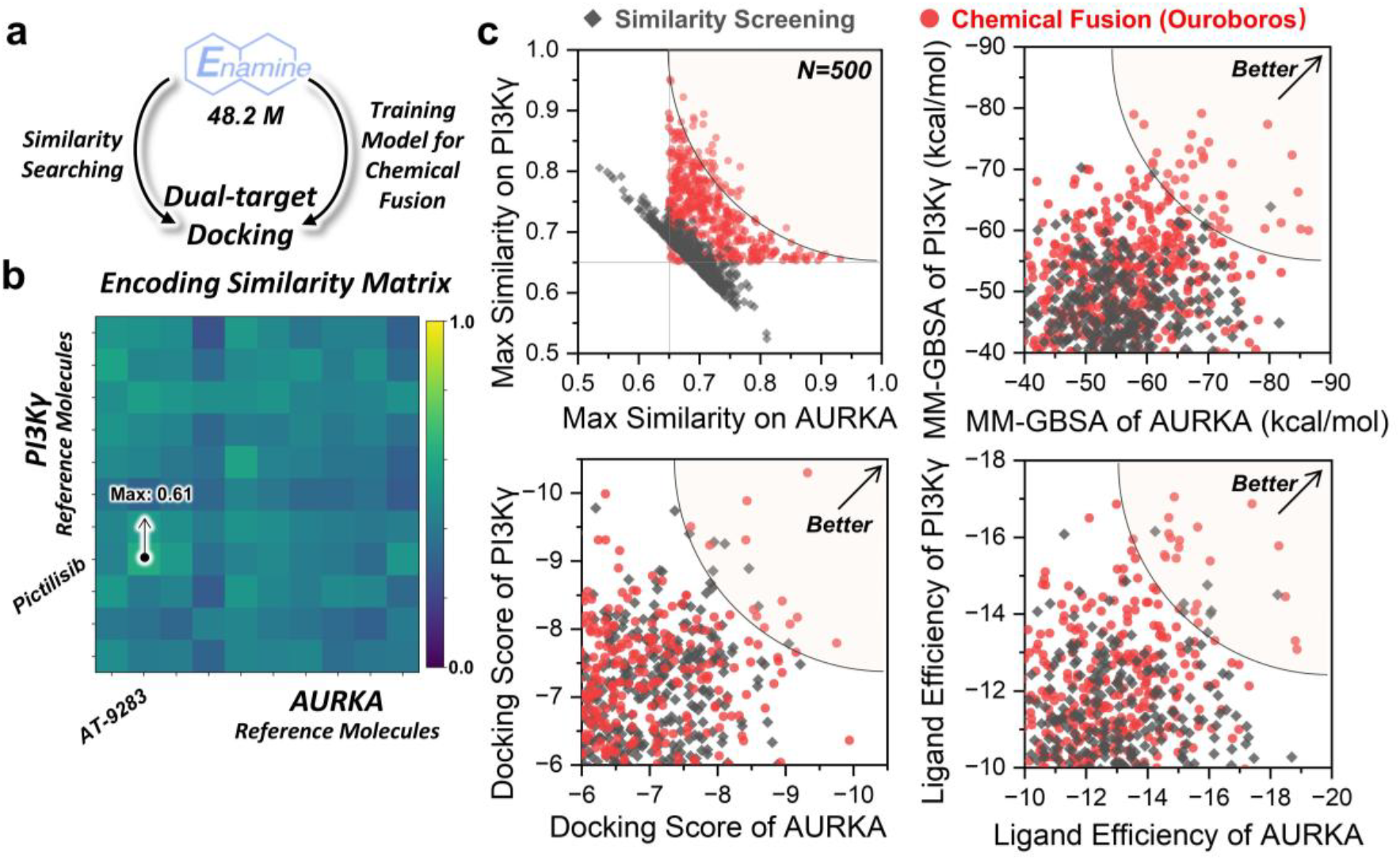
The chemical fusions in encoding space for dual-target drug discovery. **(a)** Comparison of similarity-based virtual screening and Ouroboros chemical evolution. **(b)** The encoding similarity matrix between reference molecules of AURKA and PI3Kγ. The pair of molecules with the highest similarity is pointed out with arrows. **(c)** Assessment of molecules generated using chemical fusion and similarity screening. The black diamonds represent molecules produced by similarity screening, and the red circles represent molecules produced by chemical fusion. A solid gray line in the figure represents the similarity threshold of 0.65 that was used to filter the molecules produced in the chemical fusion. The translucent sectors in the upper right corner represent high-quality candidate molecules.

#### Dual-target molecules are scarce in reference molecules

Analysis of the encoded similarity matrix for the two reference sets revealed a maximum similarity of 0.61 between them (**Figure 5b**). However, none of these compounds exhibited dual-target inhibition, suggesting that molecules with higher similarity to both reference sets might be needed to improve success rates. To address this, we retained partially fused molecules from the chemical fusion process that achieved a maximum similarity exceeding 0.65 for both targets. Among these, the top 500 molecules (ranked by synthetic accessibility) were selected and compared directly with the top 500 molecules from similarity screening in a head-to-head evaluation.

#### Chemical fusion helps design dual-target molecules

As shown in **Figure 5c**, molecules generated through chemical fusion exhibited significantly higher encoded similarity, with many achieving similarity scores above 0.7 for both targets. To further assess the development potential of these candidate compounds, we performed molecular docking to evaluate their binding affinity and calculated MM-GBSA (Molecular Mechanics Generalized Born Surface Area) binding free energy. The results demonstrated that chemical fusion produced more compounds with superior docking scores and binding free energies for both targets.

Additionally, we compared the ligand efficiency of both groups (removing the effect of ligand size), further confirming Ouroboros’ advantage in identifying highly efficient dual-target inhibitors. Overall, chemical fusion has the superior capability of lead compound screening over similarity screening, while this capability is expected to further improve as scaling up of the training dataset for molecular decoder.

## Discussion

Representation learning and generative AI have gained significant attention in drug discovery, yet a fundamental disconnect between these domains has limited the effective use of chemically meaningful representations for molecular design. To bridge this gap, we introduce Ouroboros, a novel framework that seamlessly integrates molecular representation learning with generative AI through three key components: (1) a generalized molecular encoder that projects chemical structures into 1D encoding vectors, (2) an independent structural decoder that reconstructs SMILES sequences, and (3) a chemical space navigator that enables directed molecular optimization. Notably, the decoupled training paradigm allows Ouroboros to acquire chemically relevant knowledge without generation constraints during pretraining, achieving superior performance in similarity-based virtual screening and molecular property prediction.

Ouroboros incorporates a similarity learning strategy that significantly enhances data efficiency, enabling effective pretraining on a compact chemical dataset (<150,000 molecules) while demonstrating exceptional virtual screening performance on large-scale benchmarks such as DUD-E and LIT-PCBA (>1 million molecules). Its real-world utility was further validated through multi-target drug discovery, successfully identifying three novel multi-target inhibitors from a 48.2 million-compound diversity library, showcasing its strong generalization across chemical spaces.

Architecturally, Ouroboros employs an auto-regressive Transformer decoder that reconstructs SMILES sequences from compressed 1D encodings, enabling two generative strategies: directed migration and chemical fusion. Benchmark evaluations on molecular property optimization and dual-target inhibitor design confirm its superior performance over existing methods. In this context, Ouroboros effectively bridges the gap between molecular representation learning and generative AI, establishing a new paradigm for continuous chemical evolution in encoding space. Beyond these strategies, Ouroboros can also integrate other AI-driven models, leveraging them as loss functions to optimize molecular properties.

While Ouroboros has shown superior performance in predicting multiple molecular properties, the current benchmark includes only 10 molecular properties, indicating significant opportunities for future development. Moreover, there is increasing interest in generating molecules with strong binding affinity for specific biological targets. Currently, Ouroboros does not predict drug-target binding affinity directly. Instead, it relies on molecular docking to identify molecules along the transition path with superior docking scores and binding poses. This suggests a keen need to incorporate protein representations into the training of drug-target binding affinity prediction models to further enhance the capabilities of the Ouroboros framework. With continuous progress in molecular representation learning, we anticipate that Ouroboros, supported by ongoing community-driven innovations, will evolve into a pivotal tool for AI-powered drug discovery. Its potential to streamline early-stage drug development and molecular structure optimization offers promising avenues for advancing pharmaceutical research.

## Methods

The Ouroboros protocol designs for unified drug molecule representation and generation and employs three independent modules: (1) a molecular encoder that compresses molecular structures into 1D vectors; (2) a property decoder that translates 1D vectors into molecular properties; and (3) a structural decoder that reconstructs molecular structures from 1D vectors (**Figure 1**). The protocol is flexible and can be easily extended to integrating other representation learning models with domain-specific knowledge for molecular encoding.

### AI model architecture

#### Representation and massage passing of molecular graph

To compress the chemical space, a molecular encoder of 1D molecular encoding with a size of 2048 is pre-trained by molecular similarity learning, which promotes the model to identify molecules with similar chemical structures and pharmacophore features. For this, we first converts SMILES representations into molecular graphs, which are then processed by a GNN-based Weisfeiler-Lehman Network (WLN)^31^ for features updating (**Figure S12a**). Next, a global self-attention pooling module is applied to compute atom-wise attention weights. These weights are finally used to aggregate atomic features into a molecule-level encoding vector, which can be further utilized for various pre-training tasks, including MFS and CSS.

The small molecular structure in encoder is represented as a graph *G* = (*V, E*), with *V* representing atoms and *E* chemical bonds. The atom features include atom type, hybridization, formal charge, chiral tag, whether the atom is in a ring, and whether the atom is aromatic. The bond features include whether or not they are conjugated, ring-forming, and chiral types. Both the construction of molecular graphs and the message passing are implemented through the DGL^32,33^ package.

As shown in **Figure S12a**, the GNN processes molecular graphs by iteratively updating atom and bond features through a message-passing scheme. The input atom features 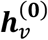 are projected into a higher-dimensional space using a linear transformation followed by a ReLU^34^ activation:

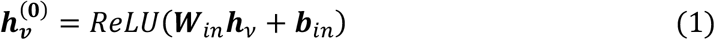

where ***W***_***in***_ is the learnable weight matrix, and ***b***_***in***_ is the bias term. For each bond feature ***e***_***u****v*_ contacting atoms ***u*** and *v*, a message is generated by concatenating the source atom features ***h***_***u***_ and ***e***_***u****v*_:

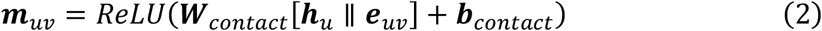

where || denotes concatenation. These messages are used to update the bonds features:

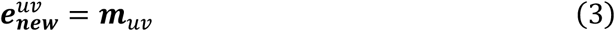

The atom features are updated by aggregating messages from neighboring bonds and combining them with the atom’s previous features:

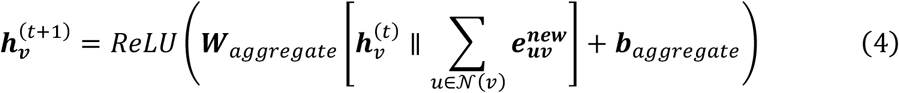

where 𝒩(*v*) is the set of neighbors of atom *v*, the *t* represents the times of massage passing (4 times in this work). Then, atom features ***h***_*v*_ and bond features ***e***_***u****v*_ are independently projected into a latent space to prepare them for further interactions, element-wise multiplicated, and aggregated from all neighbors of the atom *v*:

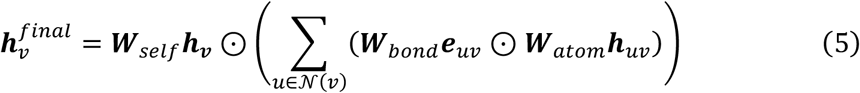

The final atom representation combines the aggregated neighbor features and a self-loop attention through element-wise multiplication.

#### Global attention pooling module

The model implements global self-attention pooling through the following steps: for each atomic features ***h***_*i*_ in the molecular graph, a MLP predicts its attention weight, denoted as *a*_*i*_. These weights are normalized using the SoftMax function to ensure they sum to 1:

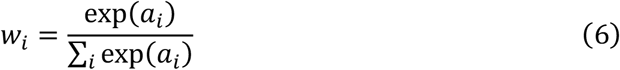

Each atomic feature is then scaled by its normalized attention weight, and the scaled features are summed to generate the graph-level encoding ***H***_*graph*_:

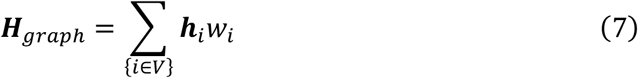

As shown in **Figure S12b**, the MLP begins by projecting the atom features from 2048 to 6144 dimensions, followed by reductions to 1024 and 128 dimensions in sequence. The linear projection in 128 dimensions is repeated three times before projecting the features to a single dimension. Batch normalization is applied after the first two linear layers, and a sigmoid activation function is applied after every linear layer, including the final layer generating the 1-dimensional attention weight.

#### Properties Decoder module for property modeling

The property predictor in Ouroboros consists of three neural network components (**Figure S12c**). The first component includes a dropout layer (dropout rate: 0.5) that projects the molecular encoding from 2,048 dimensions to 8,192 dimensions, followed by a sigmoid activation function. The second component concatenates the molecular encoding generated by the first component with the original molecular encoding, resulting in a 10,240-dimensional vector. This vector is processed through two linear layers while maintaining the feature dimension at 10,240, with LeakyReLU^35^ activation and batch normalization applied between the layers. The third component performs an element-wise addition of the input and output features from the second component. The resulting features are processed through a linear layer, batch normalization, and SiLU^36,37^ activation, then projected to 1,024 dimensions. This is followed by another round of batch normalization and SiLU^36,37^ activation. Finally, the features pass through one more linear layer with SiLU^36,37^ activation to project them to the dimensionality of the target property (1-dimensional for all properties in the benchmark).

#### Representation of chemical language

The tokenizer used in Ouroboros molecular decoder is a Byte-Pair^38^ tokenizer, in which vocabularies were obtained by segmenting the SMILES of the compounds in our dataset. These tokenizers all include at least the basic elements that make up a pharmaceutical small molecule, such as C, N, O, S, P, c, n, o, s, F, Cl, Br, I, basic chemical bonds and isomer labels, such as, “=”, “@”, “/”, “\”, and “#”, as well as other special symbols, such as brackets and numbers.

#### Molecular structural decoder module for encoding decompressor

The main role of the structural decoder module is to transform 1D encoding vectors into probability distributions with specific length tokens. It is implemented by a Transformer decoder, and the 1D encoding vector serves as the input memory. Using an autoregressive generation strategy, the decoder predicts the next token in the SMILES sequence until the end token occurred. We train a 4-layer, 32-head Transformer decoder, equipped with layer normalization and SiLU^36,37^ activation functions, to reconstruct the SMILES representation of molecular encodings by using the molecular encodings as memory tensors. For the target SMILES, rotational position encoding and an embedding layer are utilized to generate target tensors, which are used for teaching-force^39^.

A deep neural network projects hidden states onto the token vocabulary during each prediction step (**Figure S12d**). For the output features of the Transformer decoder, a 3-layer MLP is employed to project these features to match the number of tokens. Specifically, the hidden features are first transformed to 4096 dimensions, subsequently reduced to 1024 dimensions, and finally mapped to logits over 43 distinct tokens.

### Data collection and training scheme

#### Molecular datasets

We employed two molecular datasets of differing sizes to train and test the Ouroboros compressor and decompressor for molecular chemical space representation. The first dataset contains 126,248 distinct molecules, collected from five categories of resources: 1) The diverse molecular dataset previously utilized in GeminiMol^9^; 2) A macrocyclic peptide dataset CycPeptMPDB^40^; 3) The diverse molecular dataset in the Cell Painting Gallery^41,42^; 4) Molecules from the Protein Data Bank (PDB)^43^; 5) A manual-collected dataset of common biological cofactors, polypeptides, and oligonucleotides. Out of the 126,248 query molecules, 4728, 32 and 64 reference molecules are randomly selected and paired with the query molecules to form three similarity matrices of 4728×126248, 32×126248, and 64×126248, which are used as training, validation and test datasets, respectively, for the Ouroboros compression encoder. When splitting the molecule datasets, we used a Tanimoto similarity cutoff of ECFP4 < 0.3 to ensure that the molecules in the validation and test sets are non-homologous to the training set.

The second dataset is a combination of the 48.2 M Enamine REAL Diversity set^25^ and the core dataset of the 126,248 molecules, which are designed for the Ouroboros molecular decoding decompressor development. Here, the validation set comprised 200 molecular structures from the REAL dataset, 256 structures from the first four subsets in core molecular dataset, and 35 molecules in the 5^th^ subset, while all remaining molecular structures were used for training.

#### Molecular similarity matrices

We designed two types of inter-molecular similarity scores, CSS and MFS, for molecular similarity learning. All molecules in the core molecular dataset (126,248) are preprocessed, with calculations including the predictions of protonation under pH = 6.9 and tautomeric states by the LigPrep module^44^ in the Schrödinger software package. For each molecule, the protonated and tautomeric state with the lowest penalty score was selected.

For a given pair of molecules, the CSS is determined by identifying the conformations with the highest pharmacophore similarity when superimposing all conformations of one molecule onto those of the other. In our previous work on GeminiMol^9^, we implemented a simplified version of CSS descriptors, where conformational space exploration was constrained using two energy window scaling factors (0.5060 and 1.4806 kcal/mol). To generate pseudo-labels for data augmentation, we preserved the minimum similarity during similarity calculation, which was used to estimate the conformational flexibility. While this pseudo-labeling approach lacks a clear physical interpretation, the application of CSS descriptors led to improved model performance, suggesting that a more refined CSS descriptor could further enhance model accuracy.

In Ouroboros, we employ a more detailed conformational search and CSS calculation approach by incorporating four energy windows, which are defined by multiplying the number of rotatable bonds with scaling factors of 0.5060, 0.14806, 0.88836, and 1.4806 kcal/mol. Superimposition and similarity calculations are performed using PhaseShape^45^, where molecules are represented in pharmacophore space. Conformational searching is carried out using the Monte Carlo Multiple Minima (MCMM)^46^ algorithm from the MacroModel module^47,48^ within the Schrödinger software package, executing 10,000 steps under the OPLS4 force field^49^, with redundant conformations removed based on a 0.5 Å RMSD cutoff. During data augmentation, numerical calculations were streamlined by retaining only the difference between CSS values at the maximum and minimum energy windows. This strategy effectively captures molecular flexibility trends as a function of energy variations. Compared to GeminiMol^9^, the enhanced conformational search with additional energy windows, lower RMSD thresholds, and an optimized search process, is expected to improve the CSS calculation accuracy and physical interpretability without requiring complex data manipulations.

To construct the MFS matrix, three descriptors (ECFP4^50^, MACCS^51^, and AtomPairs^28^) are computed for each of the molecule pairs. For ECFP4^50^ and MACCS^51^, we employed the Tanimoto similarity measure, while for AtomPairs^28^, the Tversky similarity measure was chosen, as it demonstrated superior virtual screening performance in prior evaluations^9^. To augment the data, the difference in similarity under the largest and smallest energy windows was computed and added to the similarity vectors.

Consequently, the final similarity representation for each molecule pair contains five CSS and three MFS descriptors.

#### Similarity learning for molecular representation pre-training

The projection head of Ouroboros was designed to project the learned molecular encoding into meaningful inter-molecular similarities (**Figures S2b-c**). The projection network consists of a series of fully connected layers with non-linear activation functions and normalization. Two encoding vectors of query and reference molecules were input to the two projection heads of MFS and CSS. The encoding vector first passes through a rectifier component, which expands the features to 5 times the dimension of the original encoding vector, passes the LeakyReLU^35^ activation function and a batch normalization layer. Subsequently, the expanded features are concatenated with the original encoding vector and progressively passed through linear layers, sequentially reducing dimensions from 24,576 to 2,048, followed by batch normalization. This process continues with a reduction from 2,048 to 512 dimensions, another batch normalization step, and is repeated three times before being projected to the final output dimensions. The SiLU^36,37^ activation function was added between all linear layers in the projection head. Finally, it outputs a value that was reset to the range of 0-1 through a sigmoid neuron in the projection head.

In the pre-training task of similarity learning, we train the Ouroboros encoder by continuously sampling data from the similarity matrix (126,512 × 4,728 in training set). In each batch, there are 512 query and 48 reference molecules, with 8 different similarity labels for each pair of molecules. The model is thus trained to predict a third-order matrix (512 × 48 × 8), with the loss function being MSE between predicted and actual similarity scores.

During the similarity learning stage, training is performed using the AdamW^52^ optimizer in PyTorch^53^, with a learning rate of 5.0 × 10^−5^ and a weight decay of 0.01. Two learning rate adjustment strategies are applied: (1) gradual increase at the beginning of training, and (2) cosine learning rate scheduling, which dynamically adjusts the learning rate based on training progress to prevent convergence to local optima. In the Ouroboros encoder, the WLN parameters are inherited from the GeminiMol model and remain frozen for the first 2,000 training steps. The learning rate warmup starts at 10% of the original learning rate and increases linearly to the full learning rate over 10,000 steps. The minimum learning rate in the cosine learning rate schedule is set to 50% of the original learning rate, with a scheduling period of 10,000 steps. Additionally, an early stopping strategy is employed, where validation set performance is monitored every 200 steps. If no improvement is observed over 60 validation iterations, training is terminated early to prevent unnecessary computation.

#### Molecular Property Modeling

The training, validation, and test sets for the molecular property predictor are divided according to molecular scaffolds, ensuring that approximately 20% of the molecules in the test set possess novel scaffolds not present in the training set. During the training stage, MSE loss function and AdamW^52^ optimizer with a learning rate of 5.0 × 10^−5^ was used in training. The batch size is determined by the size of the training dataset: a batch size of 96 is used for training sets exceeding 5000 samples, 64 for those with more than 1000 samples but fewer than 5000, and 48 for training sets with fewer than 1000 samples. The learning rate scheduling strategy employed during training is identical to that used in pre-training; specifically, the period of learning rate schedule is set to 2000 steps for training sets larger than 5000 and 1000 steps for smaller training sets. Model performance is validated every 200 steps, and training is early stop if no performance improvement is observed over 60 cumulative validation evaluations.

#### Training scheme for molecular structural decoder

In the training of the molecular decoder, causal masking is applied to implement teaching-forcing^39^. To align its shape with the output tensor and integrate positional information, the teacher tensor is passed through an embedding layer followed by rotary positional encoding^54,55^. The training process uses the previously described learning rate scheduling strategy, with a period of 4000 steps. The AdamW^52^ optimizer is employed with a learning rate of 5.0 × 10^−5^ and a weight decay of 0.01.

The training loss for the molecular structure decoder is computed using a weighted cross-entropy loss. The start-of-sequence token is excluded from the loss calculation. For each token *t*_*i*_, its weight is computed as inversely proportional to its frequency *f*_*i*_ in the batch relative to the total token count, capped at 100, i.e.,

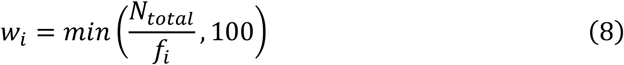

where *N*_*total*_ is the total token count in the batch. If *f*_*i*_ = 0, *w*_*i*_ is reset to 1. Padding tokens, indexed as 0 in the vocabulary, are ignored during loss computation. The token-wise cross-entropy loss (excluding padding tokens) matrix *ℒ*_CrossEntropy_ is multiplied elementwise by the weight matrix *W*:

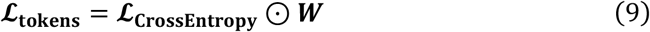

The final loss is obtained by averaging over all tokens (*j* is ground-truth token):

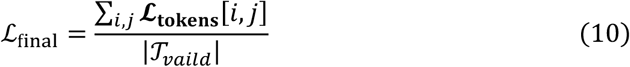

where |𝒯_*vaild*_| is the number of valid tokens (excluding start-of-sequence token).

### Downstream tasks for three independent modules

#### Virtual screening

In this study, we assessed the generalization capabilities of molecular encoders using the DUD-E^21^ and LIT-PCBA^22^ virtual screening benchmarks. In the benchmark tests of LIT-PCBA^22^, there are multiple active compounds for each target that can serve as reference molecules. Traditional baseline methods included ECFP4^50^, MACCS^51^, AtomPairs^28^, and Phase Shape^45^, while GeminiMol served as the AI-based baseline due to its use of a similar training strategy. For GeminiMol^9^, the Pearson correlation coefficient, as described in its original study, was used as the similarity metric. In the case of Ouroboros and ChemBERTa, cosine similarity was applied due to its computational simplicity, but performance comparable to the Pearson correlation coefficient. In this study, we use three different versions of ChemBERTa^26,27^, including the base model pre-trained on 100K SIMLES, and the two models Masked Language Modeling (MLM) and Multi-Task Regression (MTR) pre-trained on 77M SIMLES in ChemBERTa-2^26,27^, respectively. When using ChemBERTa to extract molecular coding, the batch size is set to 1 to avoid the effects of padding. Since the data for ChemBERTa comes from the PubChem dataset, which is the same source as the data for LIT-PCBA, we expect it to perform better.

For each query molecule *q*, we calculate the similarity to all reference molecules *R* and select the highest similarity value as the final score, i.e.,

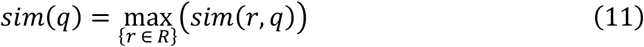

In this work, the BEDROC was used in the virtual screening as the evaluation metric, where the α of BEDROC was set to 160.9, corresponding to EF1%23.

#### Molecule property prediction

To further evaluate molecular property modeling performance, we tested various models on ten diverse drug property datasets from TDC^56^. In this study, training, validation, and test sets were split based on molecular scaffolds, following the default data partitioning settings of TDC^56^, with 20% of the data reserved for the test set. Both Ouroboros and GeminiMol^9^ use frozen molecular encoders, where fixed molecular encodings are directly projected onto molecular properties. CombineFP extracted 2048-dimensional molecular fingerprints, including ECFP4^50^, FCFP6^50^, AtomPairs^28^, and TopologicalTorsion^57^, using RDKit^58^. These fingerprints are then analyzed by AutoGluon^59^, which automated the selection of optimal machine learning algorithms from methods such as neural networks, LightGBM^60^, and CatBoost^61^. FP-GNN^24^ was tested with its default parameters and evaluated using the same data splits as the other approaches.

#### Molecular structure generation

For the structure decoder, we reconstructed molecular structures from 1D vectors in the validation set and evaluated their validity and similarity to the original molecules. For this task, the molecular datasets selected were from first four subsets of the core molecular dataset. These experiments assessed whether the model could maintain molecular validity while producing structural diversity.

We examined the model’s ability to generate diverse molecular structures by introducing 1) random Gaussian noise into 50% of the positions in the molecular encodings and 2) increasing the temperature of the autoregressive decoding process by Gumbel-max^29^ technique. This method incorporates stochasticity into the decoding process while respecting the probability distribution determined by the model.

For the first validation approach, we used Gaussian noise and scaled by *N* ϵ [0.001, 0.0025, 0.005, 0.0075]. The perturbed encoding vector ***v*** can be represented as:

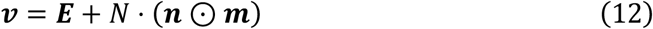

where ***n* ∼ N**(^−^**1, 1**), and for each position *i* of ***E***, the mask *m*_*i*_ is defined as:

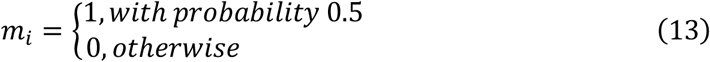

For the second approach, we used the temperatures *T* ϵ [0.01, 0.1, 0.5, 1.0, 1.5, 2.0]. To sample token indices from a predicted probability distribution, we first apply the SoftMax function with a temperature *T* > 0 to smooth or sharpen the distribution. The temperature-scaled log probabilities are computed as:

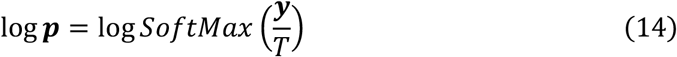

where *y* is the predicted logits, and ***p*** represents the normalized probabilities.

To enable sampling, we add Gumbel noise to the log probabilities. The Gumbel noise ***g*** is generated as:

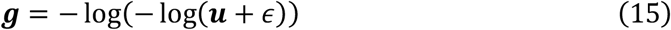

where ***u* ∼ Uniform**(**0, 1**) and *ϵ* is a small positive constant to avoid numerical instability.

The noisy log probabilities ***s*** are given by:

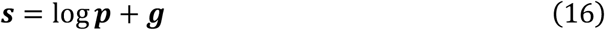

The sampled indices are obtained by taking the argmax over the ***s***.

### Applying the Ouroboros encoding in practical drug discovery

#### Biological hypothesis and reference molecules

In this study, a total of 5 cancer drivers are selected for multi-target drug screening. For KRAS, apart from itself, PI3Kα, PI3Kγ, MEK1/2, and PLK1 are selected as relevant targets. For TP53, synthetic lethal targets, WEE1 and CHK1/2, are selected. For SMAD4, AURKA is selected as a synthetic lethal target. For CDKN2A/MTAP, PRMT5 is selected as a synthetic lethal target. For BRCA1/2, PARP1/2 is selected as the synthetic lethal target. We retrieve all molecules from ChEMBL with pChEMBL values greater than 7.0 for the relevant targets. These compounds are then clustered using AtomPairs fingerprints and visually inspected to ensure structural diversity. Ultimately, 119 compounds are selected as reference molecules.

#### Similarity screening using Ouroboros encoding

In similarity screening, cosine similarity is used as the calculation metric, retaining only the maximum similarity for each reference molecule within a target set, as described in Eq. (11). For query molecules with a similarity score below 0.5, the similarity value is reset to 0. Finally, the top 0.1% of molecules, ranked by the sum of similarities, are selected for subsequent molecular docking validation.

#### Docking validation

Although Ouroboros demonstrated superior early enrichment rates in benchmark tests, however, selection of a small number of optimal candidates by careful docking inspection is necessary due to the high cost of chemical synthesis. We further enriched potential active molecules using a hierarchical virtual screening process called GVSrun (https://github.com/Wang-Lin-boop/CADD-Scripts/blob/main/GVSrun) in which the docking program is Glide^62^. The resulting docking poses were visually inspected to exclude irrational structures, which mainly include unsaturated hydrogen bond donors, ligand strain energies, and solvent effects.

#### Enzyme inhibitory assay

A total of 18 compounds are successfully synthesized and subjected to enzymatic activity assays. The selected kinases for single-dose enzymatic screening (10 µM) include PIK3CA, PIK3CG, AURKA, MEK1, CHK1, PLK1, and WEE1, while PIK3CA, PIK3CG, and AURKA are further assessed for IC_50_ determination. The enzymatic activity assay is conducted in a 384-well plate (Greiner, 784075) at 25°C for 60 minutes. Each well contains 1 μM substrate, 1 nM kinase, and 4 μM ATP in assay buffer. After the reaction, 10 μL of kinase detection reagent is added to each well, followed by incubation at 25°C, centrifugation at 1,000 rpm for 1 minute, and a final incubation at 25°C for 1 hour. Fluorescence signals are measured at 620 nm (Cryptate) and 665 nm (XL665) using a BMG plate reader. IC50 values are determined by nonlinear regression (curve fitting) using a variable slope (four-parameter model) in GraphPad Prism (v.8.0).

### Implementation of directed chemical evolution

#### Stochastic Propagation

By incrementally introducing noise to the encoding vector of the initial molecule, the progressive evolution of molecular structures can be observed. In the stochastic propagation experiment, we generate new molecular structures by adding random noise to the coding vector (**Figure S13a**).

For the entire encoding vector ***E***, we generate a standard Gaussian noise ***n* ∼ N**(^−^**1, 1**), which was scaled by the standard deviation at each location, denoted as ***σ***. The noise is further scaled to 0.05 and masked randomly at 50% of the locations. The perturbed encoding vector ***v*** is represented as:

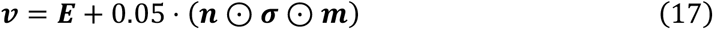

where the mask ***m*** is defined same as previous Eq. (13).

The propagation undergoes a total of 200 steps, with the molecular structure at each step generated using a molecular structure decoder operating at a temperature of 0.30. As the number of steps increases, newly generated structures are incrementally accumulated, enabling thorough exploration of the local encoding space.

#### Directed Migration

Molecular encoding vectors can be optimized using techniques similar to training neural networks, where the encoding is directionally adjusted to optimize molecular properties as defined by the loss function. Starting from the initial molecular encoding vector, the optimization is performed using the AdamW^52^ optimizer with a learning rate of 2.0 × 10^−5^ for 600 steps. After each step, the encoding vector is converted into a molecular structure using a molecular structure decoder with a temperature of 0.4 (**Figure S13b**).

The loss function consists of two terms. The first term minimizes the absolute difference between the *Sim*_*start*_ and a target value of 0.8, ensuring that structural changes are limited to preserve key pharmacophore features:

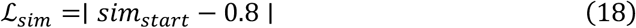

where *Sim*_*start*_ is the molecular encoding similarity between current encoding with start molecule. The second term is defined by

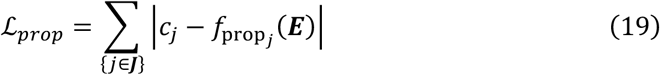

where *f*_p*ro*p_(***E***) represents the predicted molecular property, and *c*_*j*_ is the predefined target value for property *j* ϵ ***J***. This formulation guides the optimization process while maintaining a balance between multiple objectives. In this study, ***J*** consists of three molecular properties: solubility, membrane permeability, and lipophilicity.

The total loss function is expressed as:

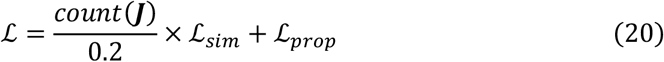

Besides optimizing molecular properties, directed migration can be used to migrate one molecular structure to another and to observe changes in molecular structure along the migration pathway within the encoding space constructed through representation learning. This method is applicable to scaffold hopping, which involves transitioning between molecular scaffolds, and to the fusion of scaffolds with two distinct properties. The loss function can be expressed as:

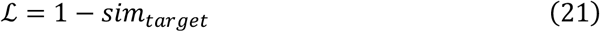

where *sim*_*target*_ is the similarity between current encoding vector and the target molecular encoding.

In the practice of scaffold hopping, the molecules generated from the directed migration are docked into the PDE4B protein structures (PDB code: 4KP6^63^ and 3O56^64^) using the Glide SP (Standard Precision)^62^ docking tool to evaluate their docking scores. The docking pose is visualized by ChimeraX^65^.

#### Chemical Fusion

Chemical fusion is an alternative approach to molecular generation by combining structural elements from reference compounds. Similar to directed migration, chemical fusion also uses optimizers and loss functions. Starting from the initial molecular encoding vector (enumerated from reference molecules), the optimization is performed using the AdamW^52^ optimizer with a learning rate 3.0 × 10^−5^ for 1,000 steps. After each step, the encoding vector is converted into a molecular structure using a molecular structure decoder with a temperature of 0.4 (**Figure S13b**).

For reference molecules *r* ϵ *R* and current molecular encoding ***E***, the similarity loss is denoted as:

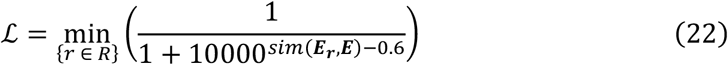

where the ***E***_*r*_ is the molecular encoding of *r*. *ℒ* could be considered as the minimum distance between the current molecule and the reference molecule, and minimizing this distance could produce a fusion between the molecules.

In the practice of multi-target drug design, the molecules generated from the chemical fusion are docked into protein structures of AURKA (PDB ID: 5DT0^66^) and PI3Kγ (PDB ID: 3ML8^67^) using the Glide SP (Standard Precision)^62^. Subsequently, the Prime module in the Schrödinger software package was used to calculate the MM-GBSA binding free energy Δ*G*_*MM*_^−^_*GBSA*_ between the docking pose and target protein, where amino acids within 5 Å around the ligand are set as flexible regions and energy minimization is practiced. Finally, the ligand efficiency *LE* is calculated by:

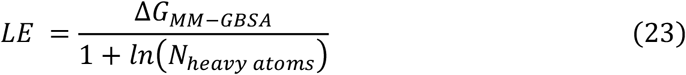

where *N*_*heavy atoms*_ refers to number of non-hydrogen atoms.

## Data availability

All molecular and benchmark datasets used in this study are collected from public resources and can be downloaded from https://zhanglab.comp.nus.edu.sg/Ouroboros/ and https://github.com/Wang-Lin-boop/Ouroboros.

## Code availability

Source codes and standalone package of the Ouroboros model are released under the MIT License and can be accessed at https://zhanglab.comp.nus.edu.sg/Ouroboros/ and https://github.com/Wang-Lin-boop/Ouroboros.

## Acknowledgements

We thank Shihang Wang, Bing Ye, Dr. Wei-kang Gong, Prof. Fang Bai, and Prof. Peng Zhan for their valuable help and insightful discussions. We also appreciate the support and services provided by ICE Bioscience Inc. in conducting the enzyme activity assays. Additionally, we are grateful to the BioHPC platform of the Institute of Systems Medicine, the Biomedical High-Performance Computing Platform of the Chinese Academy of Medical Sciences, and the Beijing Super Cloud Center for providing computational resources. This work was supported in part by the Singapore Ministry of Education (T1 251RES2309) and the National University of Singapore Startup Grants (#A-8001129-00-00 and #A-0009651-30-00). L.W. acknowledges support from the Young Scholars Program of the Chinese Academy of Medical Sciences.

## Author contributions

L.W. and Y.Z. conceived the project and designed the experiments; L.W. developed methods and designed and performed experiments; Y.W., H.L., M.L., Y.Z., C.C., C.L., and J.Z. helped with data collection and insightful discussion; L.W. wrote the initial manuscript; Y.Z. revised the manuscript; all authors proofread and approved the final manuscript.

