## Supplementary material for "Directed Chemical Evolution via Navigating Molecular Encoding Space": SI

### Supplementary Figures

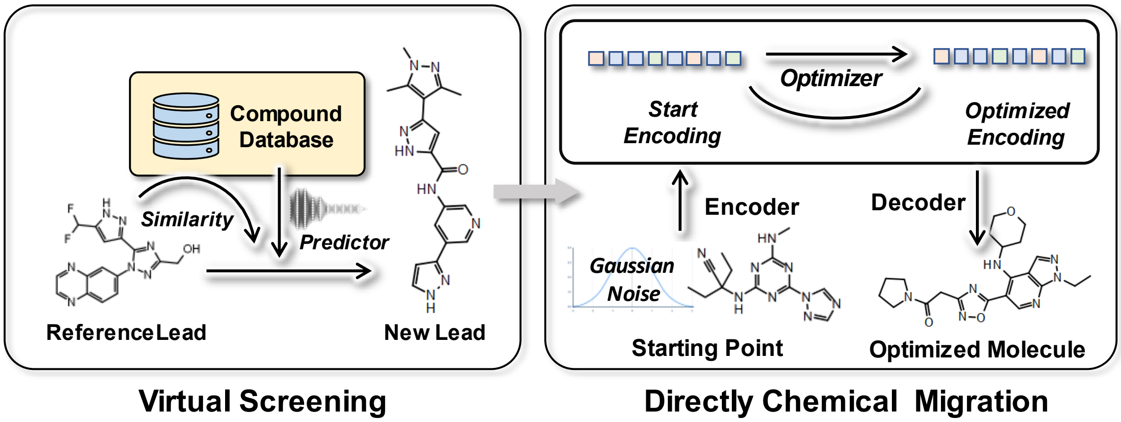


**Figure S1 | Two paradigms for designing new molecules.** First, traditional approaches to virtual screening built on representation learning models rely on the effectiveness of similarity- and predictor-based models. Second, directed molecular migration is performed within the pre-trained encoding space of representation learning. The starting point for migration can be either Gaussian noise or the encoding of an initial molecule. During the migration process, the molecular property decoder serves as a loss function to guide the direction of migration.

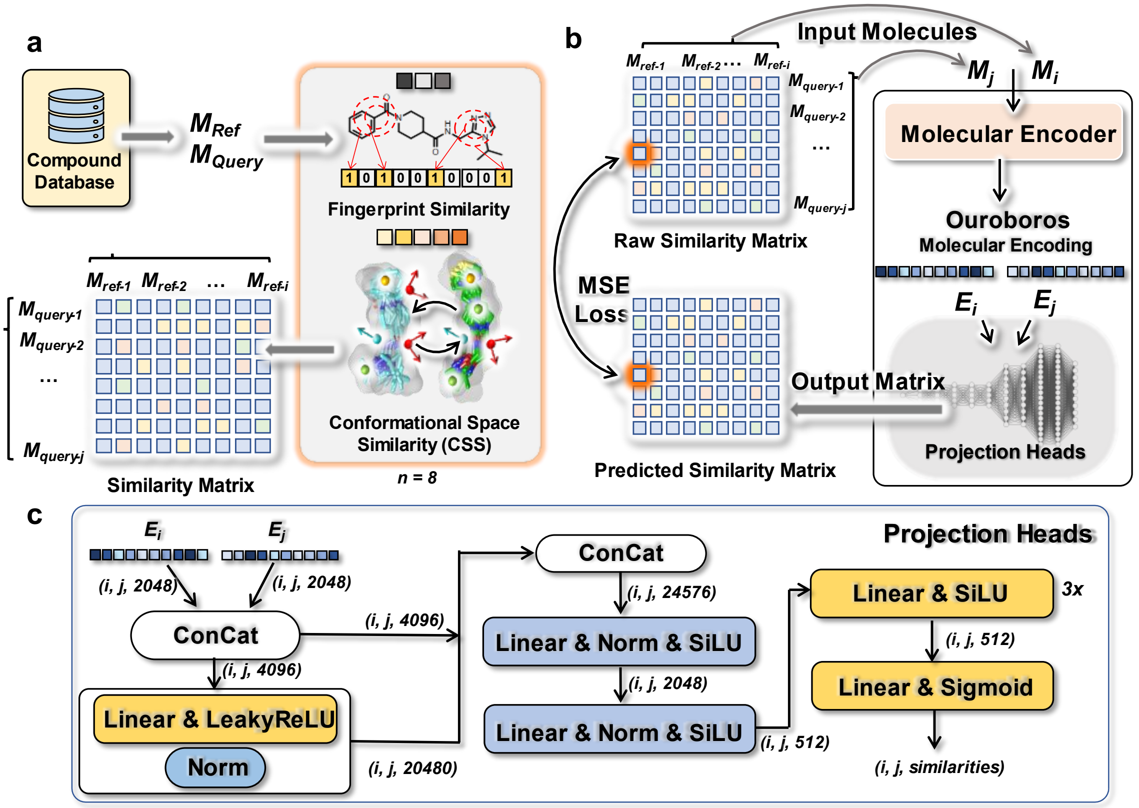


**Figure S2 | The similarity learning strategies for representation learning of Ouroboros. (a)** The construction procedure of the inter-molecular similarity dataset, where "*i*" refers to the index of reference molecule and "*j*" to index of query molecule. (**b**) Training strategies of Ouroboros encoder for compressing chemical space. (**c**) The architecture of the projection head for inter-molecular similarities.

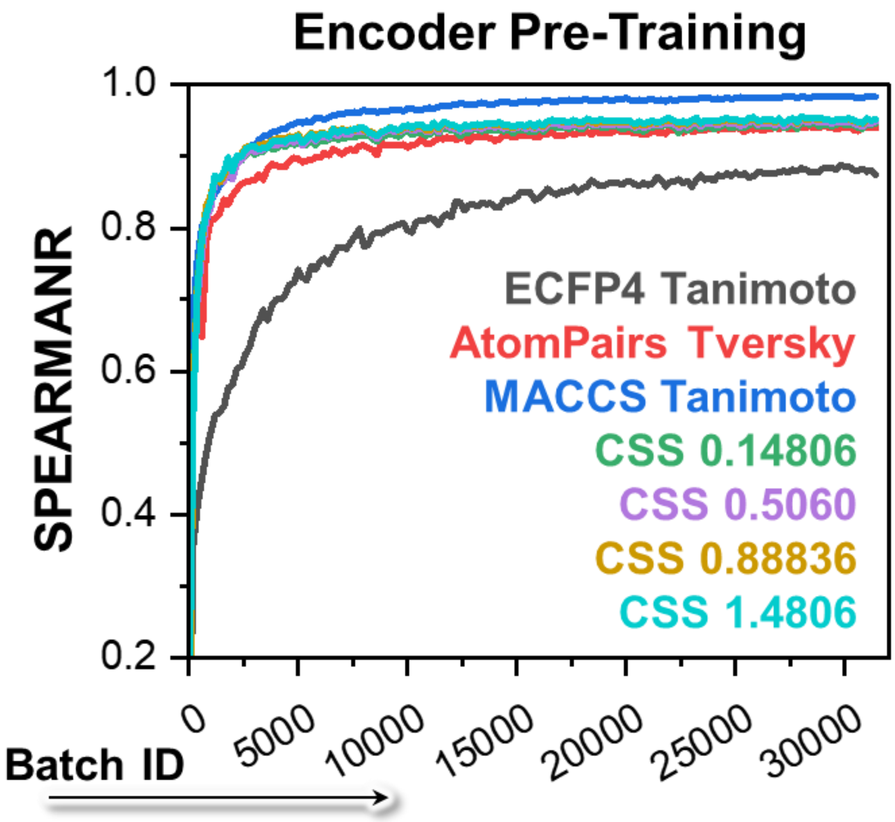


**Figure S3 | The curve of Spearman's correlation coefficient variation for the validation set during training.** The model was validated every 200 steps.

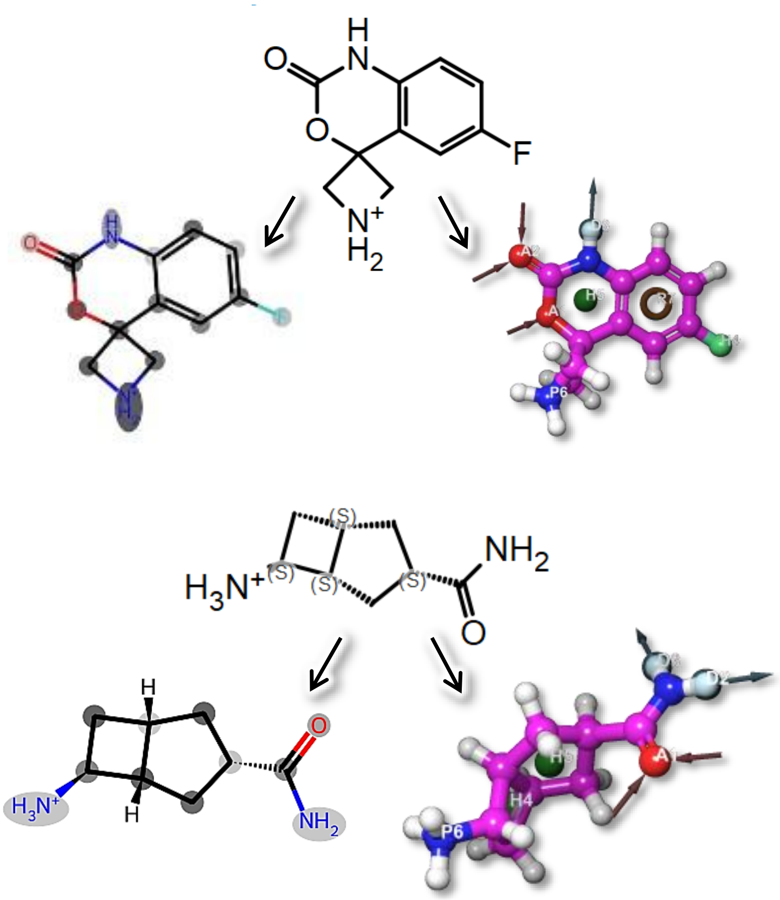


**Figure S4 | Visualization of global self-attention mechanism in molecular encoder.** In the first molecule, the nitrogen atoms in the amide and the protonated nitrogen atoms are assigned different weights, whereas in the second molecule the weights are similar, suggesting that the Ouroboros encoder is able to sense the interactions between the different functional groups within molecular structure.

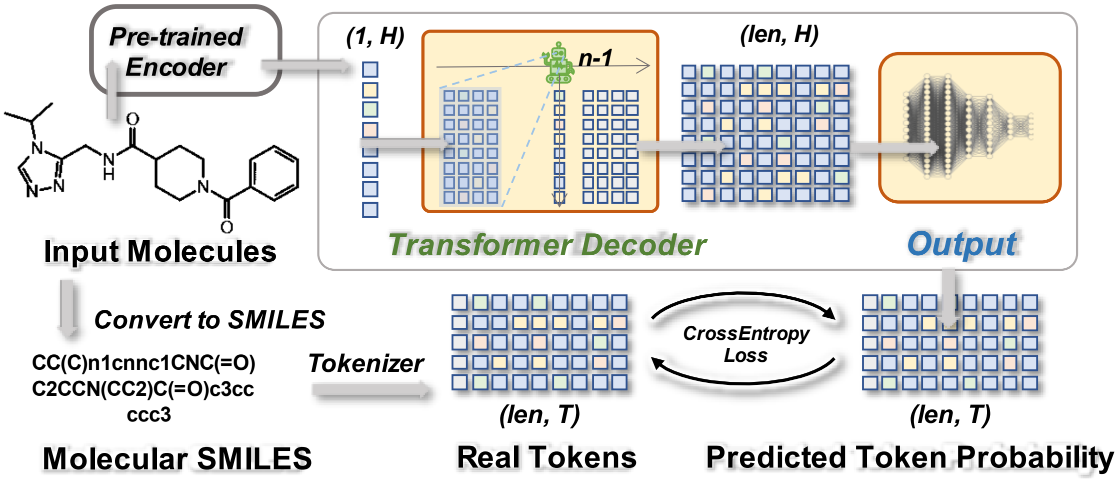


**Figure S5 | Training strategies for the Ouroboros molecular decoder.** Learnable parameters are boxed in brownish red, with 'H' referring to hidden size and 'T' to token size. The tokenization strategy for SMILES is detailed in the **Methods** section of the main text.

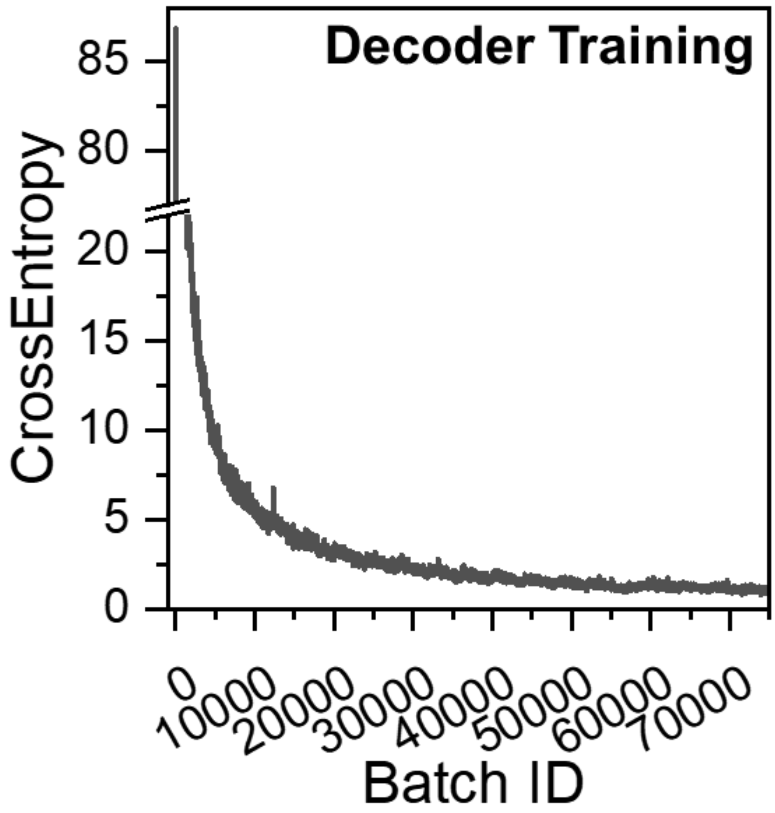


**Figure S6 | The cross-entropy loss curve during training of the molecular decoder.** The loss value recorded every 10 steps.

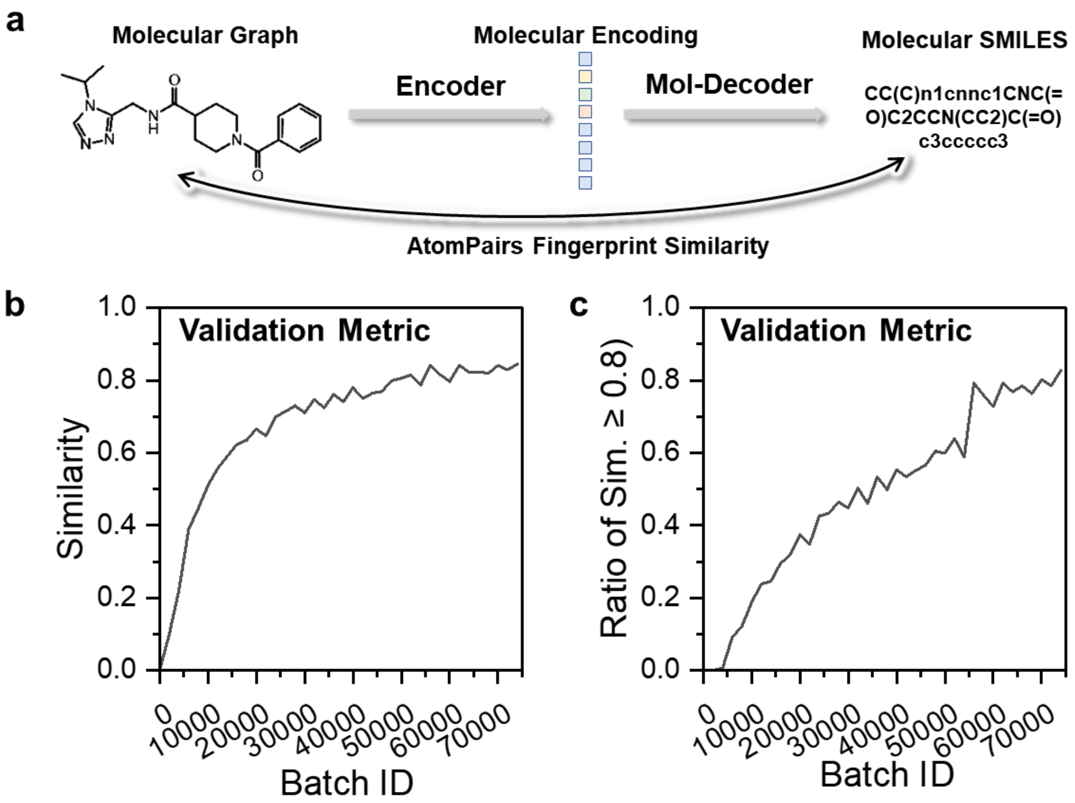


**Figure S7 | The reconstruction of molecular structure for the validation set during training stage of Ouroboros. (a)** The validation scheme for molecular decoder. The performance of the molecular decoder was evaluated by calculating the AtomPairs molecular fingerprint similarity (MFS) between the decoded and the original molecular structures. **(b)** The similarity curve of the validation set during training. The model was validated every 2000 steps. **(c)** As the model converges, the percentage of molecules that successfully recover the original structure (defined as a similarity greater than 0.8) gradually increases.

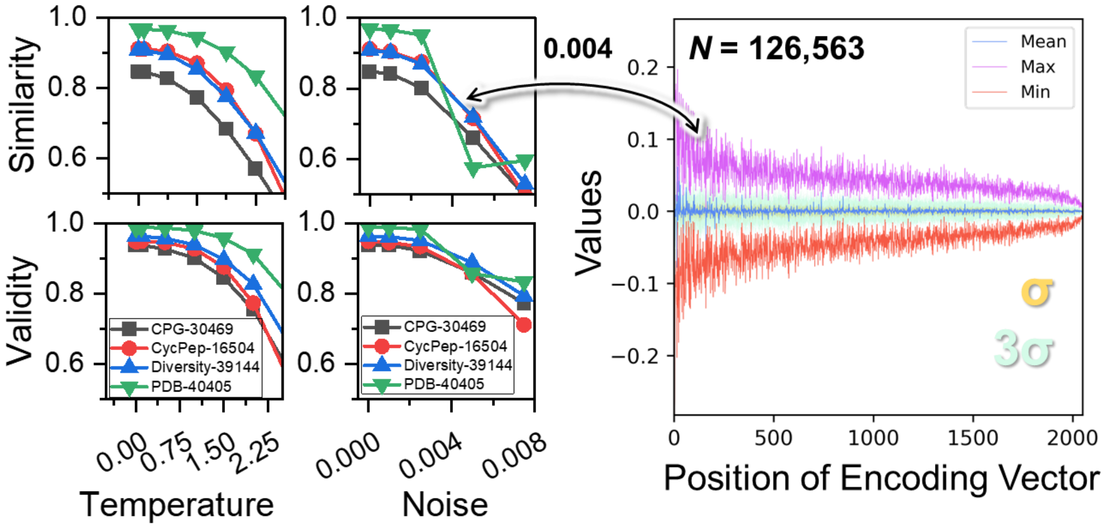


**Figure S8 | The investigations of the Ouroboros exploration capability onto neighborhood encoding space.** The temperature affects the sampling of tokens by the decoder, implemented as a Gumbel-max trick. Noise will be randomly added to 50% of the positions in the 1D encoding vector with a Gaussian distribution. The value interval of the noise is determined based on the distribution of the encoded vectors, and for the encoded vectors generated by the Ouroboros encoder, the maximum, minimum, mean, σ (standard deviation) and 3σ at each position are shown on the right. The temperature is varied in the interval between 0 and 2, and the noise is varied in the interval between 0 and 0.0075.

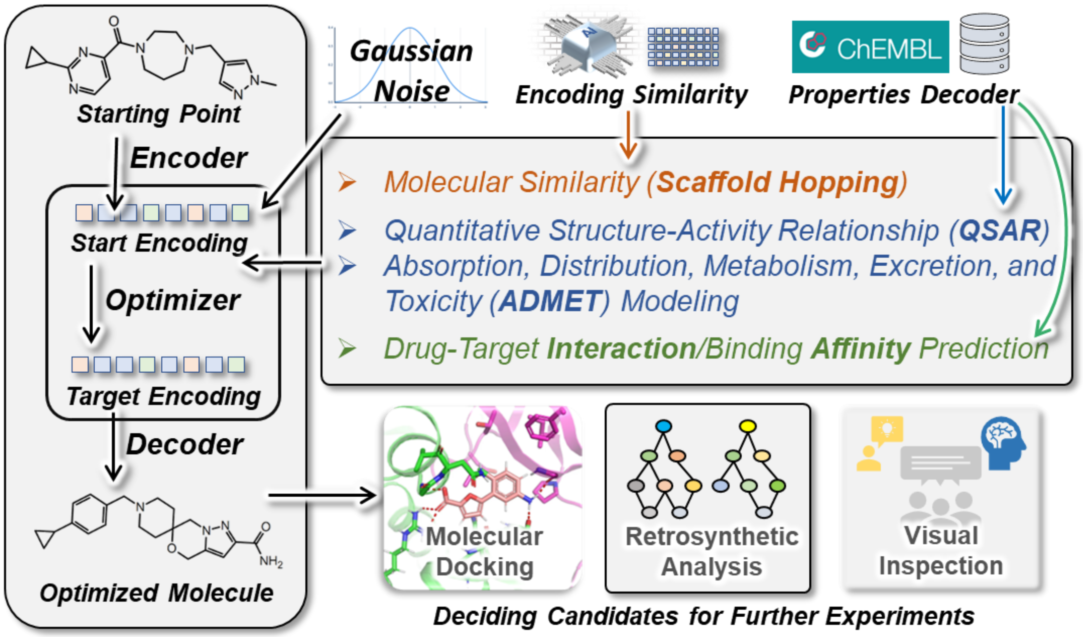


**Figure S9 | Directed chemical evolution using Ouroboros.** The encoding of the starting molecule or Gaussian noise are used to construct the starting molecular encoding, and the molecular similarity, molecular property prediction, and drug target interaction prediction are used as the loss function of the optimizer, and the molecular decoder is used to generate the molecules on the optimized pathway as outputs. Ouroboros serves as a molecular recommendation system where the produced molecules can be used for downstream molecular docking, retrosynthetic analysis and visual inspection.

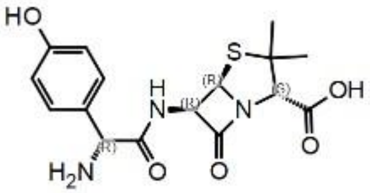

***Amoxycillin***
***Pred*** $\boldsymbol{LogP_{eff} = -5.80}$

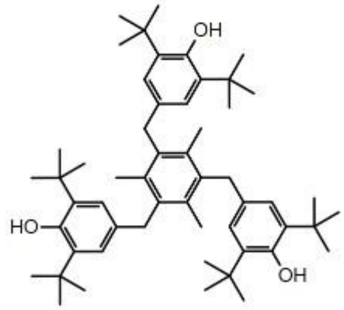

***Ionox 330***
***Pred LogS = -7.75***

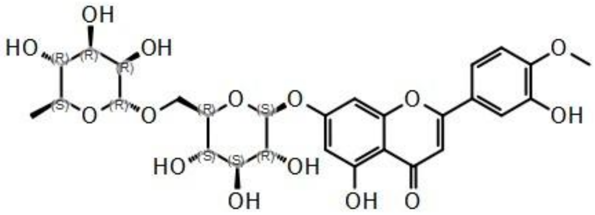

***Diosmin***
***Pred*** $\boldsymbol{LogP_{eff} = -5.88}$

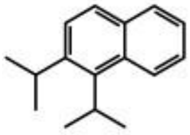

***Diisopropyl Naphthalene***
***Pred LogS = -7.29***

**Figure S10 | The organic molecules used in the benchmark of molecular property optimization.**

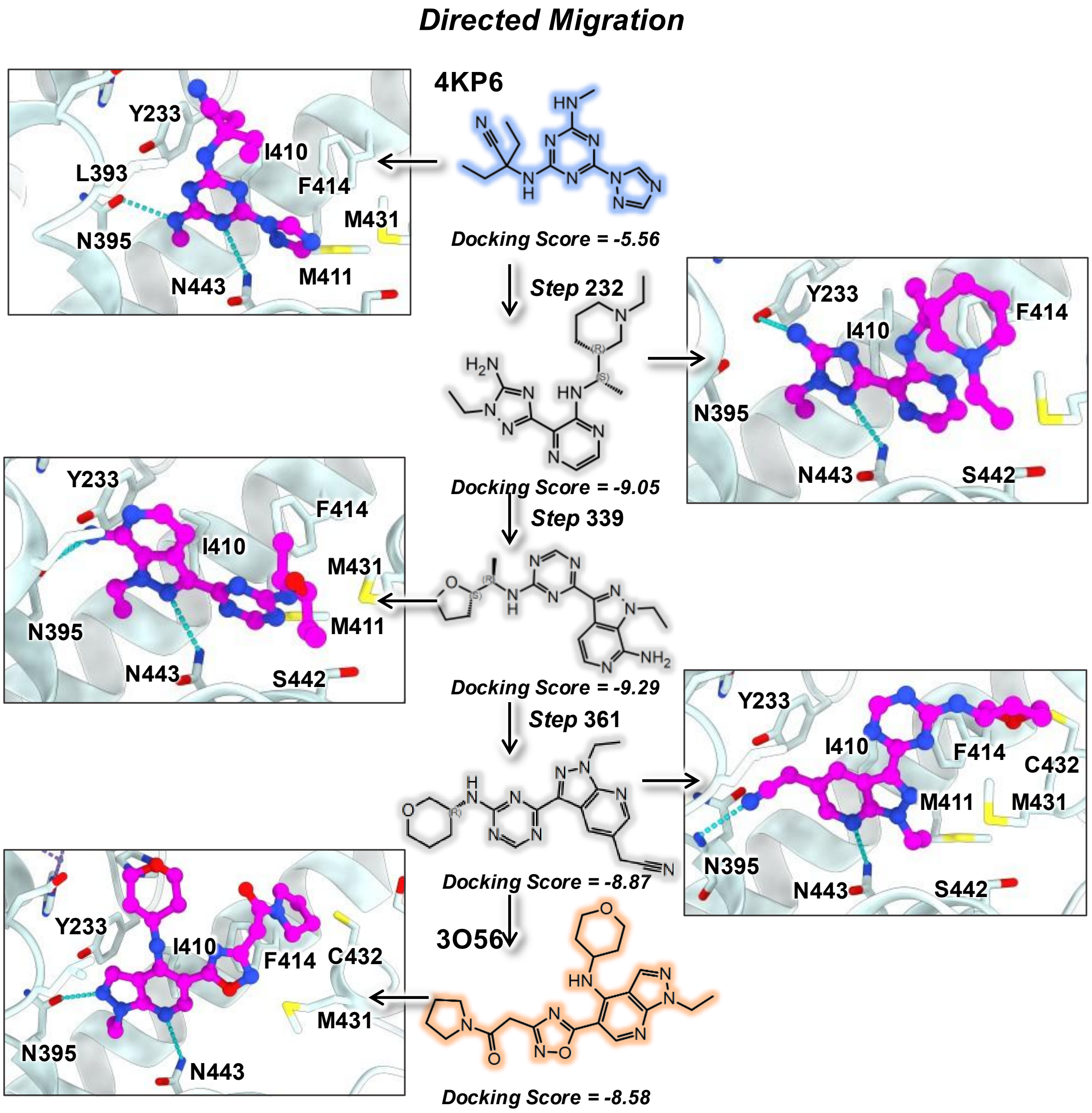


**Figure S11 | Chemical migration from one inhibitor of PDE4B to another.** Docking scores are computed using Glide SP. The C atoms of the ligands are colored in magenta.

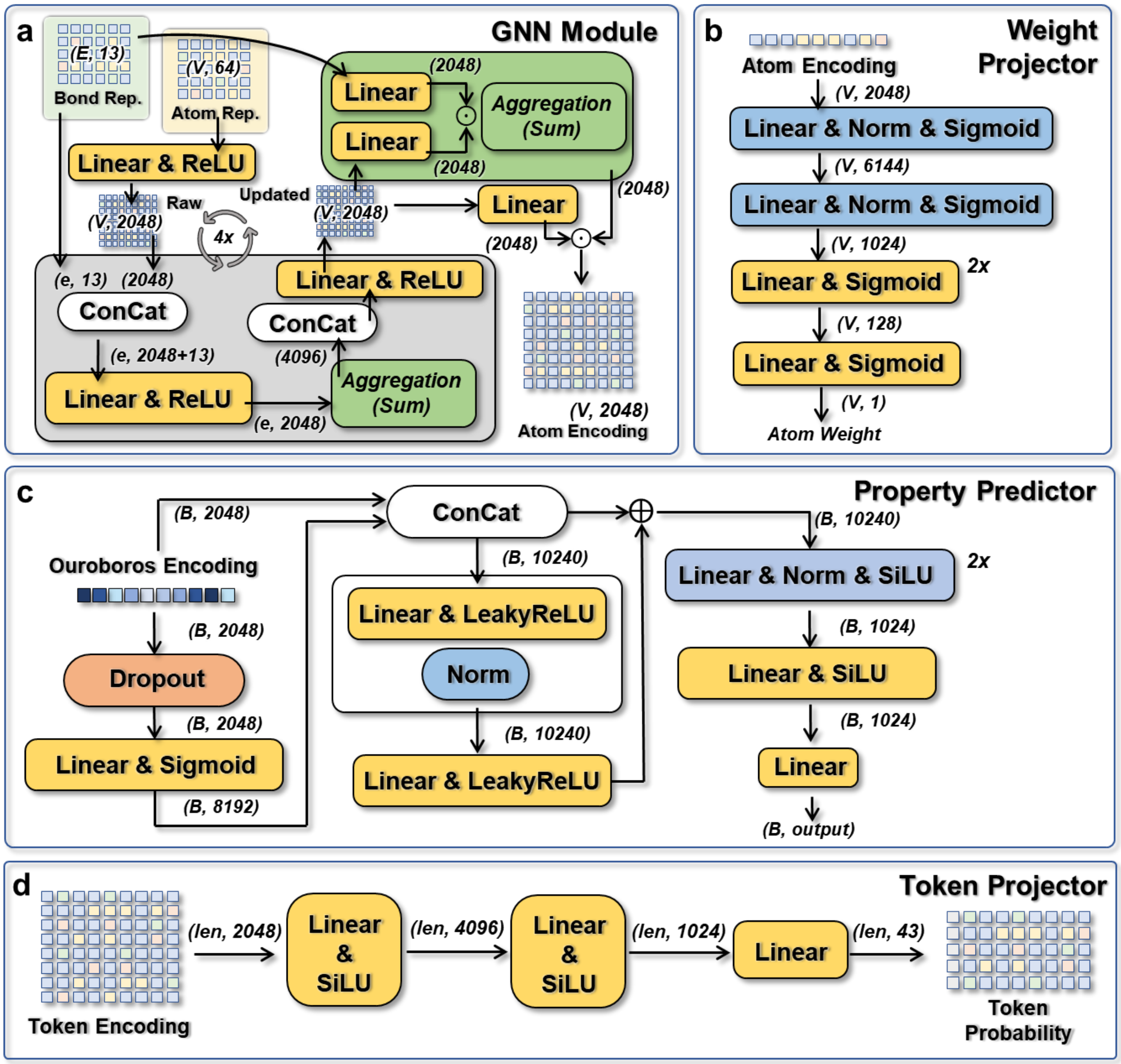


**Figure S12 | The implementation of neunal network modules in Ouroboros. (a)** GNN module for massage passing of molecular graph. **(b)** Weight projector for global attention module. Both GNN module and weight projector are used in molecular encoder, the components in the gray rounded rectangle are reused 4 times. **(c)** Property predictor for molecular proptery modeling. **(d)** Token projector was used in molecular decoder. The 'E' refers to bonds, 'V' to atoms, 'e' to bond of centor atom, 'B' to batch size, 'len' to sequence length of padded SMILES in the batch.

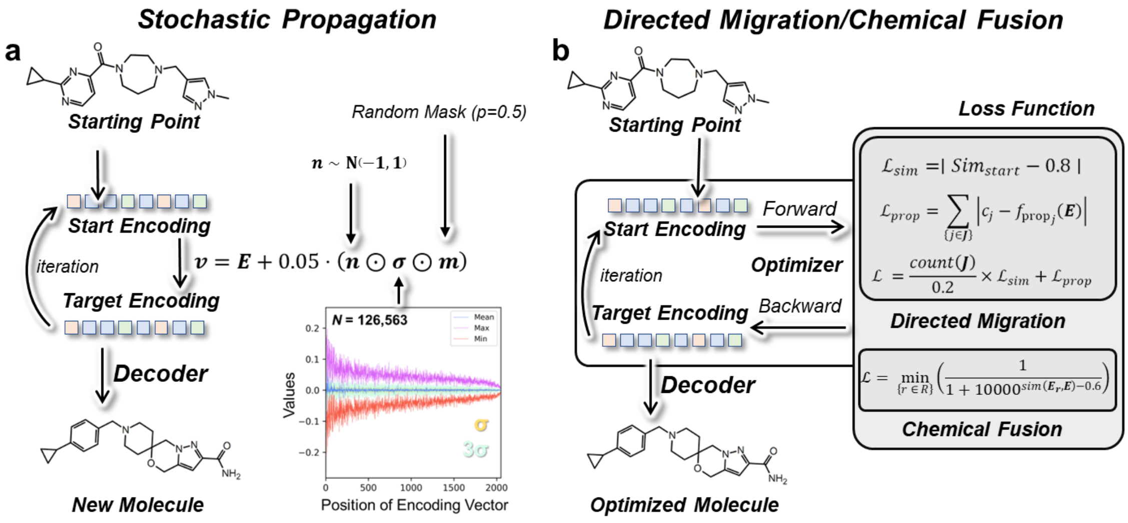


**Figure S13 | The process of stochastic propagation and directed chemical evolution. (a)** Perturbation of the compound structure is implemented by adding Gaussian noise to the encoding vector. **(b)** Directed chemical evolution by backward propagation through the optimizer. The different loss functions were used in directed migration and chemical fusion.

### Supplementary Tables

**Table S1 | Performance comparison of Ouroboros on validation and test sets in similarity learning.**

| | **RMSE** | | **PEARSONR** | | **SPEARMANR** | |
|---|---|---|---|---|---|---|
| | Validation | Test | Validation | Test | Validation | Test |
| ECFP4 Tanimoto | 0.016 | 0.016 | 0.890 | 0.878 | 0.883 | 0.874 |
| AtomPairs Tversky | 0.029 | 0.028 | 0.949 | 0.959 | 0.940 | 0.954 |
| MACCS Tanimoto | 0.023 | 0.023 | 0.983 | 0.979 | 0.983 | 0.977 |
| CSS_0.14806 | 0.046 | 0.035 | 0.933 | 0.953 | 0.946 | 0.955 |
| CSS_0.5060 | 0.044 | 0.033 | 0.938 | 0.958 | 0.950 | 0.960 |
| CSS_0.88836 | 0.043 | 0.033 | 0.940 | 0.960 | 0.954 | 0.962 |
| CSS_1.4806 | 0.043 | 0.033 | 0.941 | 0.960 | 0.955 | 0.962 |

**Table S2 | Virtual screening benchmark results on DUD-E and LIT-PCBA.**

| **Methods** | **AUPRC** | **AUROC** | **BEDROC[1]** | **EF0.01%** | **EF0.1%** | **EF1%** | **logAUC** |
|---|---|---|---|---|---|---|---|
| *Performance on DUD-E* | | | | | | | |
| Ouroboros | **0.363** | 0.777 | **0.557** | 32.177 | 29.925 | 20.272 | **0.472** |
| GeminiMol | 0.307 | 0.734 | 0.490 | 30.704 | 27.227 | 17.718 | 0.418 |
| ECFP4 | 0.345 | 0.753 | 0.556 | **33.196** | **30.230** | **20.360** | 0.450 |
| AtomPairs | 0.278 | 0.752 | 0.469 | 32.319 | 27.656 | 16.469 | 0.408 |
| MACCS | 0.230 | 0.720 | 0.380 | 29.306 | 22.792 | 13.289 | 0.365 |
| ChemBERTa[2] | 0.202 | 0.679 | 0.351 | 27.932 | 22.079 | 11.969 | 0.330 |
| ChemMLM[3] | 0.295 | **0.782** | 0.476 | 30.721 | 26.681 | 16.961 | 0.431 |
| ChemMTR[3] | 0.239 | 0.686 | 0.415 | 30.352 | 24.030 | 14.678 | 0.363 |
| *Performance on LIT-PCBA* | | | | | | | |
| Ouroboros | **0.030** | 0.579 | **0.078** | 110.511 | **24.925** | **7.575** | **0.222** |
| GeminiMol | 0.029 | **0.592** | 0.074 | 112.501 | 24.022 | 6.559 | **0.222** |
| ECFP4 | 0.021 | 0.527 | 0.053 | 70.127 | 16.361 | 4.483 | 0.189 |
| AtomPairs | 0.024 | 0.561 | 0.058 | 66.269 | 22.368 | 5.809 | 0.206 |
| MACCS | 0.017 | 0.557 | 0.043 | 32.764 | 10.752 | 3.891 | 0.196 |
| ChemBERTa[2] | 0.015 | 0.575 | 0.042 | 61.445 | 15.910 | 4.771 | 0.197 |
| ChemMLM[3] | 0.024 | 0.579 | 0.058 | **112.685** | 17.698 | 6.038 | 0.207 |
| ChemMTR[3] | 0.021 | 0.508 | 0.043 | 99.056 | 19.073 | 4.672 | 0.175 |

[*] Performance for the best model for each metric are bolded.
[1] BEDROC is calculated under the α set to 160.9.
[2] ChemBERTa-zinc-base-v1, pre-trained on 100 K SMILES from ZINC.
[3] ChemBERTa-77M, MLM and MTR, pre-trained on 77 M SMILES from PubChem.

**Table S3 | Spearman correlation coefficients on the property modelling test sets**

| Property | Size | Ouroboros | CombineFP | GeminiMol | FP-GNN |
|---|---|---|---|---|---|
| Caco2 | 818 | 0.791 | 0.778 | 0.755 | **0.843** |
| Clearance Hepatocyte | 1091 | **0.500** | 0.416 | 0.465 | 0.399 |
| Clearance Microsome | 992 | **0.601** | 0.586 | 0.546 | 0.557 |
| Half Life | 601 | **0.537** | 0.333 | 0.355 | 0.338 |
| Hydration Free Energy | 587 | **0.886** | 0.797 | 0.876 | 0.845 |
| LD50 | 6641 | 0.635 | **0.651** | 0.541 | 0.585 |
| Lipophilicity | 3780 | **0.816** | 0.747 | 0.733 | 0.790 |
| PPBR | 1453 | 0.557 | 0.510 | 0.620 | **0.637** |
| Solubility | 8876 | 0.833 | 0.823 | 0.802 | **0.835** |
| Steady-State Distribution Volume | 1017 | **0.540** | 0.472 | 0.509 | 0.300 |
| Mean | / | **0.670** | 0.611 | 0.620 | 0.613 |

* Performance for the best model for each task are bolded.

**Table S4 | Inhibitory activity of 18 candidate compounds on 7 kinase targets.**

| Compound ID # | AVE Inhibition% (10 µM) | | | | | | |
|---|---|---|---|---|---|---|---|
| | PIK3CA | PIK3CG | AURKA | MEK1 | CHK1 | PLK1 | WEE1 |
| 1 | 19.03 | -2.64 | 27.94 | -7.70 | 3.66 | -7.29 | 2.51 |
| 2 | **60.18** | 14.77 | **73.57** | -12.02 | 4.87 | **54.49** | 15.76 |
| 3 | 11.86 | 30.81 | **67.10** | -4.12 | 3.26 | 3.49 | 5.67 |
| 4 | 19.05 | 17.62 | 31.44 | -5.99 | 10.08 | -12.88 | 8.13 |
| 5 | 9.16 | -2.68 | 31.20 | -1.95 | 8.08 | -1.30 | 7.67 |
| 6 | **60.71** | 29.76 | **87.25** | 10.14 | 25.38 | -18.68 | 0.18 |
| 7 | 2.88 | -2.08 | 22.31 | -6.28 | -0.69 | -6.89 | 5.05 |
| 8 | 10.18 | 8.05 | **58.96** | -7.64 | 1.48 | -15.37 | 8.87 |
| 9 | 20.71 | 19.79 | **52.94** | 0.06 | -0.49 | -5.46 | 5.92 |
| 10 | 11.42 | 13.51 | 40.26 | -20.06 | -2.24 | -22.99 | 6.84 |
| 11 | 36.66 | 23.65 | **83.30** | 12.20 | 3.54 | -4.19 | 12.91 |
| 12 | 9.99 | 4.92 | 38.74 | -21.91 | -5.82 | -17.73 | -0.15 |
| 13 | **52.82** | **86.07** | **96.00** | 14.02 | 4.20 | 4.02 | 44.00 |
| 14 | 7.28 | 4.28 | 23.75 | -3.36 | 7.02 | -2.35 | -1.34 |
| 15 | 38.79 | 16.36 | 38.56 | -3.02 | -1.46 | 41.26 | 5.63 |
| 16 | 42.96 | 32.00 | 29.45 | 13.65 | 3.47 | 9.62 | 7.97 |
| 17 | -1.05 | 4.28 | 32.46 | -6.02 | 0.04 | -7.03 | 7.42 |
| 18 | 23.76 | 9.31 | 12.50 | -4.09 | 1.98 | -7.67 | 2.87 |

* All data are repeated twice and the values in the table are averaged. Values with inhibitions above 50 % are colored and bolded in black.
